# A novel repressor-activator-competitor module comprising C_2_H_2_ zinc finger and NAC transcription factors regulates rice grain development

**DOI:** 10.1101/2024.07.15.603577

**Authors:** Priya Jaiswal, Richa Priyadarshini, Antima Yadav, P V Aswathi, Arunima Mahto, Iny Elizebeth Mathew, Upasana Das, Falah Qasim, Ankur Vichitra, Akanksha Panwar, Ankit Verma, Akhilesh K. Tyagi, Pinky Agarwal

## Abstract

Grain size and quality are crucial agronomic traits. We have characterized a seed-preferential C_2_H_2_ zinc finger transcriptional repressor, *ZOS1-15*. Its overexpression, knock-down and knock-out plants indicated a negative control over grain size due to altered cell expansion. ZOS1-15 homodimerized and directly interacted with co-repressor TOPLESS and histone deacetylases to form a repression complex. ZOS1-15 also interacted with Mediator subunit MED14_1 and a seed-preferential transcriptional activator, ONAC024, with three alternatively spliced isoforms. The ectopic expression of *ONAC024* negatively affected plant growth and development. Seed-preferential overexpression and knock-down plants showed ONAC024 as a positive regulator of grain length due to increased cell proliferation and expansion. CRES-T generated transgenic rice plants indicated a functional divergence amongst ONAC024 isoforms. Tandem interactions were observed between ONAC024-ONAC023-ONAC026-ONAC020. ZOS1-15 and ONAC024 functioned antagonistically to regulate grain amylose and SSP accumulation while ONAC023 affected only amylose. ZOS1-15 and ONAC024 directly regulated the expression of two SSP encoding genes. Binding of ONAC024 was competed by ONAC025-MADS29 complex. The seed-preferential overexpression of SS1/ ONAC025 resulted in decreased grain size and amylose content, but higher yield. This study proposes a ’repressor-activator-competitor’ module, wherein ZOS1-15, ONAC024, ONAC023, ONAC025 along with their interactors synergistically and antagonistically regulate multiple aspects of rice grain development.

## INTRODUCTION

Rice is a valuable staple food for more than half of the world’s population with grain as the commercially important part. Grain yield and quality are the two foremost components influencing rice production. Grain yield depends on the number of panicles per plant, number of grains per plant and grain size. Grain size is a multifaced character regulated by multiples genes. It depends on three parameters, grain length, width and weight. So far, several QTLs/genes regulating numerous traits of rice seed development have been documented. Some of the important genes controlling grain yield are, *GW5* (Weng et al., 2008), *GW8* (Wang et al., 2012), *GW7* (Wang et al., 2015), *DPE1* (Dong et al., 2015), *GS3* (Nan et al., 2018), *GW6* (Shi et al., 2020), *GW2* (Verma et al., 2020), *RBG1* (Lo et al., 2020), *LARGE2-APO1/2* (Huang et al., 2021), *OsPDCD5* (Dong et al., 2021), *WD40* (Chen et al., 2022) etc. Grain quality is evaluated by four major criteria: nutrition, appearance, milling, cooking and eating texture (Hori and Sun, 2022). In general, rice grain quality and size are closely associated with its development. Rice grain development is a continuous process that involves several molecular and cellular events. Seed development is divided into three landmark events: cell division, organ initiation and maturation covering five stages (S1-S5) (Agarwal et al., 2011). This whole process of seed development is ruled by an intricate network of transcription factors (TFs) working in a spatio-temporal coordination.

TFs are DNA binding proteins that play vital roles in controlling gene regulatory networks by modulating the expression of downstream genes. Several plant TF families such as zinc finger proteins (ZFPs), MYB, bHLH, bZIP, WRKY, AP2-ERF, HD-ZIP, NF-Y and NACs play important roles in regulating diverse developmental processes, including seed development (Mathew and Agarwal, 2018; Das et al., 2019; Duan et al., 2021). ZFPs form a diverse and large group of plant TF families. ZFPs are classified into several types such as C_2_H_2_, C_2_HC, C_2_HC_5_, C_3_HC_4_, CCCH, C_4_, C_4_HC_3_, C_6_ and C_8_ based on the number of cysteine and histidine residues bound to the zinc ion (Schumann et al., 2007). Amongst all classes of ZFPs, C_2_H_2_ TFs are one of the largest classes of ZFPs. C_2_H_2_ ZFP was first discovered in *Petunia* as *EPF1*, which possesses plant-specific QALGGH motif in the zinc finger domain (Takatsuji et al., 1992). In rice, a total of 189 C_2_H_2_ ZFPs have been identified and named as ZOS (Zinc finger of *Oryza sativa*) (Agarwal et al., 2007). C_2_H_2_ ZFPs play a wide variety of functions in regulating plant developmental processes such as spikelet development in rice (Zhuang et al., 2020), GA biosynthesis (Duan et al., 2021), flower development (Lyu and Cao, 2018), pollen and seed development (Han et al., 2018; Puentes-Romero et al., 2022) and vegetative and reproductive development (Xu et al., 2018; Liu et al., 2022c).

Gene regulation is inextricably linked to the transcriptional state of chromatin which can be either activated or repressed. Transcriptional repressors are proteins that bind to native transcription sites and repress gene expression (Reynolds et al., 2013). Many plant C_2_H_2_ ZFPs harbor an active repression motif. They repress target gene expression by interacting with a co-repressor and a chromatin remodeling factor (Ohtani and Iwasaki, 2021, Nishioka et al., 2020, Xun et al., 2022). The predominant repression motif present in various TFs families like AUX/IAA, MYB, C_2_H_2_ and WRKY are ethylene responsive element binding factor associated amphiphilic repression (EAR) motif (Plant et al., 2021). EAR motif is represented by the consensus sequence DLNxxP and LxLxLx where x can be any amino acid (Singh et al., 2019). EAR motif containing repressors are known to interact with co-repressors such as TOPLESS (TPL), TOPLESS RELATED (TPR), SIN3A ASSOCIATED PROTEIN18 (SAP18) as well as members of the histone deacetylase (HDAC) family and together the complex regulates multiple developmental processes (Yang et al., 2018; Hu et al., 2020; Deng et al., 2022).

Another large group of plant-specific TFs are the NACs (NAM, ATAF1/2, CUC2). There are 151 NAC TFs in rice (Nuruzzaman et al., 2010). The protein consists of a typical conserved NAC domain at the N-terminal and a diversified C-terminal transcriptional regulatory region (TRR) (Christianson et al., 2010). The NAC domain is additionally divided into five sub-domains from A-E. It has been reported that sub-domains C and D are highly conserved and have the DNA binding sites (Mathew and Agarwal, 2018). The TRR domain is liable for transcriptional activation or repression properties (Zhao et al., 2016). NACs function in a wide array of plant growth and development processes, such as floral whorl boundary formation, xylem vessel differentiation, embryogenesis, secondary wall thickening, leaf development, seed germination, lateral root development, fruit development and senescence (Laubscher et al., 2018; Mathew and Agarwal, 2018; Trupkin et al., 2019; Jia et al., 2021; Ma et al., 2021; Kou et al., 2021; Liu et al., 2022a; Singh et al., 2021; Vargas-Hernández et al., 2022). Few NACs have been identified for their role in seed development such as OsNAC129 (Jin et al., 2022); NARS1 and NAC2 (Kunieda et al., 2008); NAM-B1 (Waters et al., 2009) and ONAC025/ SSI (Mathew et al., 2020). NACs usually function as a homodimer or heterodimer protein for stable DNA binding (Harrington et al., 2019) such as with MADS TF (Cordenunsi-Lysenko et al., 2019; Zhang et al., 2021), GSK2 and SAPK8 (Wu et al., 2022), CrMYB68 (Zhu et al., 2020). The dimer OsNAC20-OsNAC26 regulates starch and seed storage protein synthesis (Mathew et al., 2016; Wang et al., 2020b), ONAC127-ONAC129 regulates rice grain filling (Ren et al., 2021), ZmNAC128-ZmNAC130 dimer transcriptionally regulates the accumulation of starch and protein in seeds (Zhang et al., 2019) and OsCUC2-OsCUC3 dimer regulates organ boundary formation (Wang et al., 2021).

This study establishes the role of a repressor-activator-competitor module consisting of C_2_H_2_ ZF TF, ZOS1-15, as the repressive component, ONAC024 as a strong activator and SS1/ ONAC025 as the competitor. The components of the repressor, activator and competitor complex have been elucidated. Detailed phenotypic and biochemical analysis of transgenic and/ or mutant plants of members of this module, *ZOS1-15, ONAC024*, *ONAC023* and *SS1/ ONAC025*, have been generated and analyzed to determine their regulation of grain length, width, weight, yield, amylose and protein contents. Further, the *cis* elements to which ZOS1-15, ONAC024 and SS1/ ONAC025 bind to, in the promoters of two downstream genes, and their combinatorial regulation, have been established. Our study establishes the existence of a C_2_H_2_ ZF-NAC TF module which regulates multiple aspects of rice grain development.

## RESULTS

### ZOS1-15 negatively regulates plant architecture, grain size and flowering time in rice

To investigate the role of *ZOS1-15*, ectopic overexpression (1-15_OE), knock-down (1-15_KD) and knock-out (1-15_KO) rice transgenic plants were raised in *indica* rice genotype, IET-10364 (**Figure 1A**). High expression levels of *ZOS1-15* in 1-15_OE (197-459 folds) and downregulation (39.3-73.3 folds) was seen in 1-15_KD transgenic grains with respect to wild type (**Supplemental Figure 1**). 1-15_KO rice plants were raised targeting either the ZF domain (ZF1-15) or the C-terminal of DLN motif sequence (sig1-15) (**Supplemental Figure 2A, B, C**). For ZF1-15 construct, transformed calli did not survive on the regeneration media, and turned black (**Supplemental Figure 2D**). Thus, complete disruption of ZOS1-15 indicated lethality. For sig1-15 construct, a total of 11 transgenic lines were obtained, out of which 10 lines showed mutation (**Supplemental Figure 2E**). In sig1-15 CRISPR knock-out plants, Sanger sequencing data revealed that three lines showed a 3 bp GGA deletion (named 1_1-15_KO) which was homozygous. Four lines had a heterozygous monoallelic genotype with a 3-bp GGA deletion and an unmutated copy (2_1-15_KO). Another line showed a T insertion (3_1-15_KO). Two lines had a ‘T’ insertion and a 3-bp GGA deletion (4_1-15_KO). To examine the inheritance of targeted mutations in the next generation, the target region from the next generation of two lines (1_1-15_KO and 4_1-15_KO) was sequenced. Both lines had a 3-bp GGA deletion and were homozygous (**Supplemental Figure 2F**). No editing was observed in either of the predicted off-target genes (**Supplemental Figure 3**).

**Figure 1.**
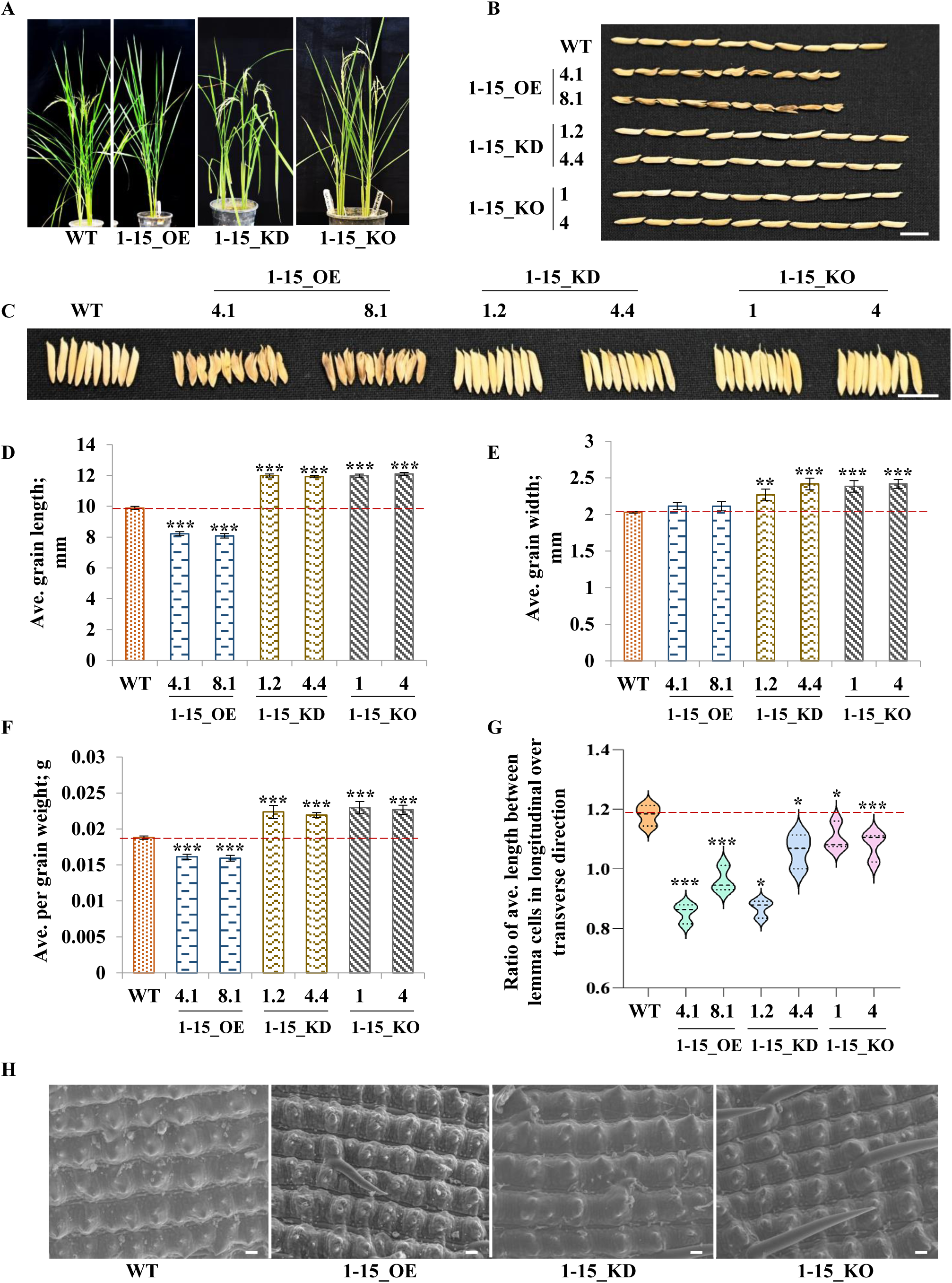
*ZOS1-15* negatively regulates grain size. **(A)** Phenotype of mature rice plants of WT (wild type), with overexpression construct of *ZOS1-15* (1-15_OE), RNAi construct of *ZOS1-15* (1-15_KD) and CRISPR mutants of *ZOS1-15* (1-15_KO). Ten mature grains for two lines each of 1-15_OE (4.1 and 8.1; T4 grains), 1-15_KD (1.2 and 4.4; T4 grains) and 1-15_KO (1 and 4; T3 grains) and WT have been arranged **(B)** lengthwise and **(C)** widthwise, scale bar = 1 cm. Bar graphs show **(D)** average per grain length, **(E)** average per grain width and **(F)** average per grain weight for two lines each of 1-15_OE, 1-15_KD, 1-15_KO and WT, n=50*3. The data was calculated from three independent plants for each line. **(G)** Violin plot shows ratio of average length between lemma cells in longitudinal over transverse direction in 1-15_OE, 1-15_KD and 1-15_KO grains as compared with WT, n=45 (15*3). Asterisks denote significant difference as determined by Student’s t-test (**\***,**\*\***,**\*\*\*** = p value ≤ 0.05, ≤ 0.01, ≤ 0.005, respectively). Error bars = ± SE **(H)** SEM photographs of outer surface of lemma of mature grains from 1-15_OE, 1-15_KD and 1-15_KO along with WT, scale bar = 20 μm.

Grain phenotyping of 1-15_OE transgenic plants showed a decrease (16-18%) in grain length (**Figure 1B, D**). Concomitantly, grain length was increased in 1-15_KD and 1-15_KO (21-22%) transgenic grains. Scanning electron microscopy (SEM) of grain husk (**Figure 1G-H**, **Supplemental Figure 4)** showed that the lemma cell number was unchanged in 1-15_OE, 1-15_KD and 1-15_KO grains. The lemma cell length was decreased in 1-15_OE and increased in 1-15_KD and 1-15_KO grains **(Supplemental Figure 4A, B)**. Hence, ZOS1-15 negatively affected grain length by hindering cell expansion. Further, 1-15_OE grains did not show change in grain width **(Figure 1C, E)** while an increase was observed in 1-15_KD (11-18%) and 1-15_KO (17-18%) grains. In transverse direction, number of lemma cells again remained unchanged **(Supplemental Figure 4C, D)** in transgenic grains with respect to WT, but lemma cell width was decreased in 1-15_OE and increased in 1-15_KD and 1-15_KO, indicating that ZOS1-15 negatively affected grain width by influencing cell expansion. Additionally, ratio of the average length between lemma cells in longitudinal over the transverse direction was lowest in 1-15_OE grains (**Figure 1G**). Hence, ZOS1-15 negatively regulates cell expansion in both orientations. 1-15_OE had decreased grain weight (15-16%) (**Figure 1F**). There was an increase in grain weight in 1-15_KD (26-28%) and 1-15_KO (∼29%). In addition, the OE grains had an open beak phenotype (**Figure 1B, C**). The results implied that ZOS1-15 negatively regulated grain length, width and weight. Though a complete knock-out of ZOS1-15 was detrimental for plant formation, a knock-down of ZOS1-15 or a knock-out targeting DLN motif was beneficial for grain size.

In addition to grain size, ZOS1-15 affects plant height, leaf morphology and panicle number. The plant height of 1-15_OE plants was reduced while 1-15_KD and 1-15_KO plants were taller (**Figure 1A**, **Supplemental Figure 5A**). The stem thickness in 1-15_KD and 1-15_KO was reduced and was unaffected in 1-15_OE plants (**Supplemental Figure 5B**). Flag leaf length and width, second leaf length, and panicle number were reduced in all plants (**Supplemental Figure 5C-F)**. 1-15_OE plants had shorter panicles (**Supplemental Figure 6A-B**). Grain filling was highly reduced, and unfilled grains were increased in 1-15_OE, 1-15_KD and 1-15_KO plants, which resulted in reduced grain yield (**Supplemental Figure 6C-E**). Further, 1-15_OE had delayed flowering **(Supplemental Figure 7A)**. The relative expression of flowering time related genes (*OsNF-YB9, OsNF-YC12, OsMADS50, OsDTH8, OsHd1* and *OsHd3a)* in 56 days old leaves was increased while *OsRFT1* expression was decreased in 1-15_OE plants **(Supplemental Figure 7B)**.

### ZOS1-15 forms a functional complex with other regulators

TFs including C_2_H_2_ ZFs often dimerize in order to execute their function (McCarty et al., 2003; Lu et al., 2010). ZOS1-15 homodimerized as shown by yeast two-hybrid, bimolecular fluorescence complementation (BiFC) and pull-down assays. To determine the region responsible for interactions, three deletion constructs of ZOS1-15 were made (**Figure 2A**). Growth of yeast cells co-transformed with ZOS1-15_AD and ZOS1-15_BD on SD/-AHLT media and blue color on SD/-AHLT + X-α-gal plates indicated homodimerization (**Figure 2B**; **Supplemental Figure 8A**), for which the presence of both ZF domains was essential. DLN motif was not needed for homodimerization. BiFC assay showed that ZOS1-15 dimer associated with cell membrane (**Figure 2D**). Further, this interaction was confirmed by pull down assay (**Figure 2E**) where the presence of 44.92 kDa band of GST_ZOS1-15 was visible when MBP_ZOS1-15 was bound to beads and the 61.42 kDa band of MBP_ZOS1-15 was observed when GST_ZOS1-15 was bound to beads. *In planta* validation of homodimerization of ZOS1-15 in *Nicotiana benthamiana* leaves by co-immunoprecipitation assay (Co-IP) revealed that YFP_ZOS1-15 (46.81 kDa) was coimmunoprecipitated with ZOS1-15_MYC (**Figure 2F**).

**Figure 2.**
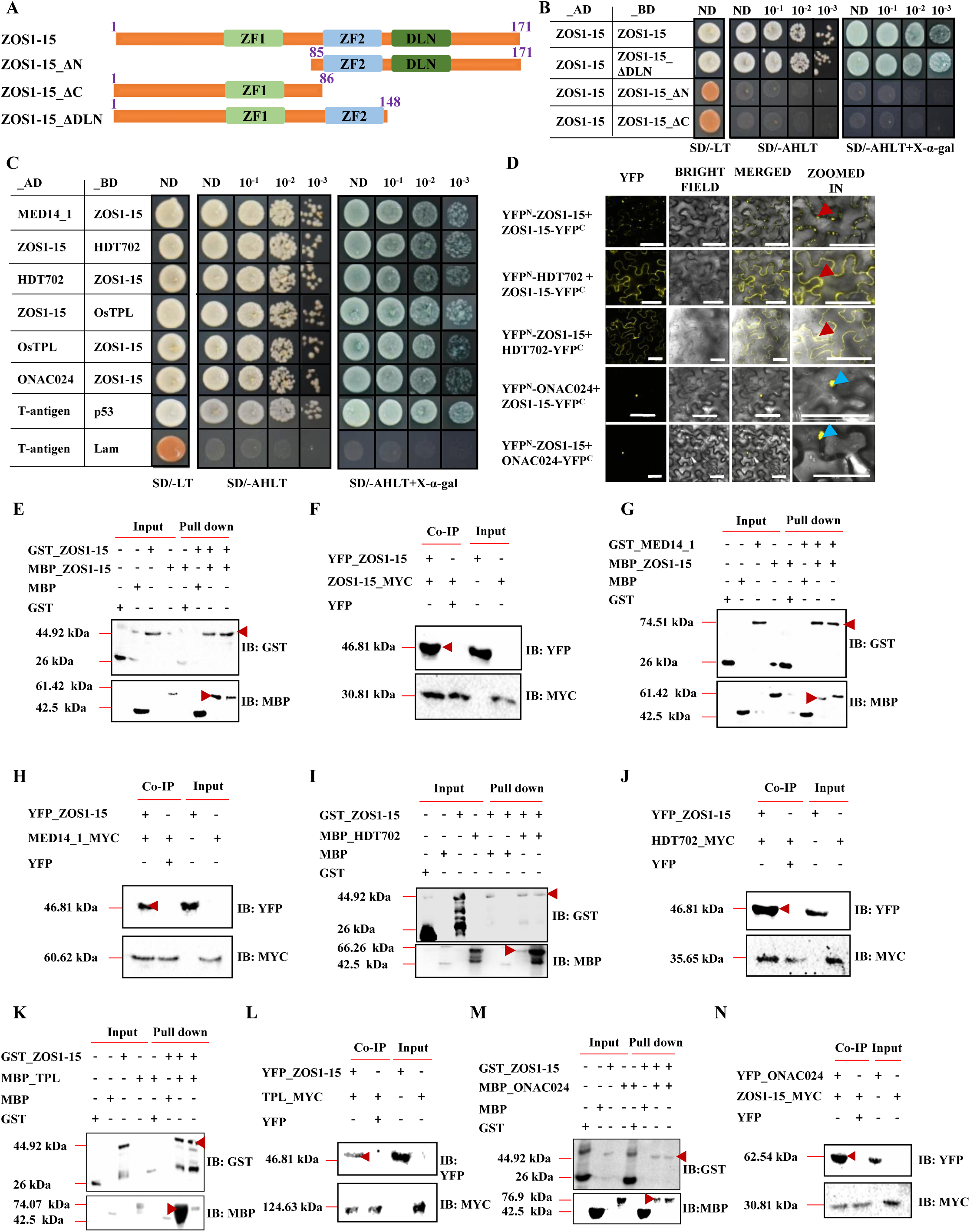
ZOS1-15 homodimerizes and forms a repression complex with MED14_1, HDT702 and TPL; and interacts with ONAC024. **(A)** Schematic representation of ZOS1-15 and its truncated forms. ZF1 and ZF2 represent C_2_H_2_ zinc fingers, DLN represents the repression motif, ΔN, ΔC and ΔDLN represent deletions of the respective regions. Diagram is not drawn to scale. **(B)** Y2H assays show homodimerization of ZOS1-15 and **(C)** its interacting partners, MED14_1, HDT702, OsTPL and ONAC024. _AD and _BD represent fusion with GAL4 activation domain or binding domain, respectively. SD/-LT indicates synthetic drop-out medium lacking leucine and tryptophan. SD/-AHLT indicates media lacking adenine, histidine, leucine, tryptophan and SD/-AHLT + X-α-gal indicates SD/-AHLT plus 5-bromo-4-chloro-3-indolyl-α-D-galactopyranoside (X-α-gal). p53_BD and T-antigen_AD are positive controls while Lam_BD and T-antigen_AD are negative controls. ND, 10^-1^, 10^-2^, 10^-3^ denote no dilution, 10-fold dilution, 100-fold dilution and 1000-fold dilution respectively, of co-transformed yeast cultures. **(D)** BiFC assays show subcellular localisation of dimers of ZOS1-15 and its interactors in *Nicotiana benthamiana* leaf cells. YFP^N^ and YFP^C^ indicate N-terminal and C-terminal regions of YFP in fusion with the mentioned protein. Plasma membrane has been indicated by red arrow and nucleus is indicated by blue arrow. Left panel indicates reconstituted YFP signal at 514 nm. Merged indicate the merger of YFP and bright field and the right column shows the zoomed-in image of the interactions. Scale bar is 50 μm. Pull down and co-immunoprecipitation assays, respectively, confirm **(E-F)** homodimerisation of ZOS1-15 [GST_ZOS1-15 (44.92 kDa), YFP_ZOS1-15 (46.81 kDa)] and its interaction with **(G-H)** MED14_1 [GST_MED14_1 (74.51 kDa), YFP_ZOS1-15 (46.81 kDa)]; **(I-J)** HDT702 [MBP_HDT702 (66.26 kDa), GST_ZOS1-15 (44.92 kDa), YFP_ZOS1-15 (46.81 kDa)]; **(K-L)** TPL [MBP_OsTPL (74.07 kDa), GST_ZOS1-15 (44.92 kDa), YFP_ZOS1-15 (46.81 kDa)]; **(M-N)** ONAC024 [GST_ZOS1-15 (44.92 kDa), YFP_ONAC024 (62.54 kDa)]. Input, pull down fractions and the antibody used have been marked. Pull down and Co-IP lane with expected protein bands have been indicated by red arrow. IB denotes immunoblot.

Mediator subunits interact with specific TFs controlling various processes (Malik et al., 2020). ZOS1-15 interacted with Mediator 14_1 (MED14_1) by yeast two-hybrid assay (**Figure 2C**). For this, full length ZOS1-15 was essential (**Supplemental Figure 8B, C**). The presence of 74.51 kDa band of GST_MED14_1 when MBP_ZOS1-15 was bound to beads and the 61.42 - kDa band of MBP_ZOS1-15 was observed when GST_MED14_1 was bound to beads in the pull-down assay confirmed the formation of this dimer (**Figure 2G**). An *in vivo* Co-IP assay confirmed this interaction by the detection of YFP_ZOS1-15 (46.81 kDa) on the immunoblot (**Figure 2H**). ZOS1-15 protein has a repression motif DLNYPP at the C-terminal (Singh et al., 2019). Repressor proteins recruit co-repressors and HDACs to form repression complexes (Cheng et al., 2018). ZOS1-15 interacted with members from all three classes of HDACs, including HDT701, HDT702, HD704 and SRT701 (**Figure 2C**, **Supplemental Figure 8D**). ZOS1-15 also interacted with OsTPL in yeast two-hybrid assay (**Figure 2C**). The complete ZOS1-15 protein was needed for interaction with OsTPL and HDT702 (**Supplemental Figure 8B, C**). BiFC assay showed the association of ZOS1-15-HDT702 dimer with the cell membrane (**Figure 2D**). The presence of 44.92 kDa band of GST_ZOS1-15 when MBP_HDT702 was bound to beads and the detection of 66.26 kDa band of MBP_HDT702 when GST_ZOS1-15 was in bound fraction, confirmed the dimer in a pull-down assay (**Figure 2I**). *In planta* Co-IP assay in *Nicotiana benthamiana* leaves further validated this interaction by the detection of YFP_ZOS1-15 (46.81 kDa) (**Figure 2J**). Interaction of ZOS1-15 with OsTPL was confirmed by the presence of 74.07 kDa band for MBP_OsTPL when GST_ZOS1-15 was in bound fraction and the detection of 44.92 kDa band for GST_ZOS1-15 when MBP_OsTPL was bound to beads (**Figure 2K**). The interaction between ZOS1-15 and TPL was further confirmed by *in planta* Co-IP assay and YFP_ZOS1-15 (46.81 kDa) band was detected on the blot (**Figure 2L**).

ZF TFs often interact with other TFs (Tran et al., 2007). ZOS1-15 interacted with seed-preferential ONAC024 as shown by yeast two-hybrid assay (**Figure 2C**). This dimer was formed in the presence of both ZFs; and the DLN motif was not necessary for this interaction (**Supplemental Figure 8B, C**). ONAC024 has the structure of a typical NAC TF (**Supplemental Figure 12A**). The TRR (transcriptional regulatory region) domain of ONAC024 was responsible for this interaction **(Supplemental Figure 9).** BiFC assay showed the localization of the dimer to the nucleus (**Figure 2D**). In the pull-down assay, the presence of 44.92 kDa band of GST_ZOS1-15 when MBP_ONAC024 was bound to beads and the 76.9 kDa band of MBP_ONAC024 when GST_ZOS1-15 was bound to beads, confirmed the formation of the ZOS1-15-ONAC024 dimer (**Figure 2M**). Further, ZOS1-15-ONAC024 dimer was validated by *in planta* Co-IP assay by the detection of YFP_ONAC024 (62.54 kDa) on the blot (**Figure 2N**). Both Y2H and BiFC results indicated that the homodimerization and interaction of ZOS1-15 with ONAC024 was distinct and did not require DLN motif as compared with its interaction with other regulators, MED14_1, HDT702 and OsTPL.

### Both ZOS1-15 and ONAC024 are seed-preferential transcription factors with highly regulated localization

Microarray data has indicated a preferential and high expression of both *ZOS1-15* and *ONAC024* in different seed developmental stages (Sharma et al., 2012). This was confirmed by qRT-PCR **(Supplemental Figure 10A)**. The highest expression was seen in S4 stage (850-fold) for ZOS1-15 and S3 stage (208-fold) for ONAC024. Both genes were differentially expressed at low levels in various other tissues like panicle (P2), internode, node, flag leaf and leaf sheath. In S1-S3 stages, the expression of *ONAC024* was higher than *ZOS1-15* while it was reversed in S4-S5 stages.

ZOS1-15 had putative nucleolar retention signal (NoRS), nuclear localization signal (NLS) and nuclear export signal (NES) (**Supplemental Figure 10B**). In YFP tagged sub-cellular localization experiments of full length ZOS1-15 and its deletion constructs, in onion peel **(Supplemental Figure 10B, C)** and *N. benthamiana* leaf epidermal cells (**Supplemental Figure 11A)**, ZOS1-15 faintly localized to the nucleus and was present in punctate structures throughout the cell. These structures partially overlapped with the ER marker and did not overlap with Golgi body and plastid markers. The localization of YFP-(ΔNoRS)-1-15 was similar to full length ZOS1-15, except in the nucleolus, indicating the presence of a NoRS at the N-terminal. YFP-(ΔNES)-1-15 localized to both nucleus and nucleolus, confirming NoRS at N-terminal and NES at C-terminal. YFP-(ΔNoRSΔNES)-1-15 localized to nucleus and cytoplasm and no punctate structures were formed. This confirmed the presence of a NLS.

ONAC024 was 309 residues long with the typical structure of a NAC TF **(Supplemental Figure 12A**). *ONAC024* showed the existence of different isoforms with non-canonical 5’ donor and 3’ acceptor sites resulting in the formation of three different transcripts **(Supplemental Figure 12D)**. Sequence analysis of alternatively spliced transcripts confirmed **(Supplemental Figure 12B**) this as short direct repeat (SDR) mediated splicing (Niu et al., 2010). A small region of the third exon was being spliced out. Semi-quantitative RT-PCR (**Supplemental Figure 12C, D**) showed that the full-length transcript was the main form and was named as ONAC024_A. The next abundant forms were named as ONAC024_B (by splicing out of 85 bp) and ONAC024_C (39 bp spliced out). Confirmation of alternative splicing was done by Western blotting (**Supplemental Figure 12E**). YFP:ONAC024 expression in *Nicotiana benthamiana* epidermal leaf cells showed two distinct protein bands of 62.54 kDa (for ONAC024_A) and 57.76 kDa (for ONAC024_B) in the immunoblot. A thick band at 62.54 kDa probably indicated a merger of ONAC024_A with ONAC024_C (60.98 kDa). ONAC024 was a transcriptional activator **(Supplemental Figure 13A)** with ONAC024_A as the strongest activator, followed by ONAC024_C. ONAC024_B with a codon shift in TRR, could not activate reporter genes. Deletion analysis showed that TRR was responsible for the strong transcriptional activation property of ONAC024 **(Supplemental Figure 13B).** The dimerization of isoforms of ONAC024_B and ONAC024_C with ZOS1-15 was weaker as compared with ONAC024_A **(Supplemental Figure 9A)**, with decreasing levels of interaction observed with ONAC024_C and ONAC024_B, respectively.

All three isoforms of ONAC024 (YFP-ONAC024_A, YFP-ONAC024_B and YFP-ONAC024_C) were localized to the cytoplasm and stayed outside the nucleus in onion peel (**Supplemental Figure 10D, E**) and *N. benthamiana* leaf epidermal cells (**Supplemental Figure 11B)**. It can be concluded that ZOS1-15 has a highly regulated localization pattern while ONAC024 requires an interacting partner for nuclear localization. Further analysis in this article elaborates on this module of a repressor-activator composed of ZOS1-15 and ONAC024.

### *ONAC024* positively regulates grain size

To understand the functional role of *ONAC024*, it was ectopically expressed in rice (024_OE) using the maize *UBIQUITIN* promoter. A total of 16 transgenic lines were analysed (**Supplemental Figure 14, 15A**). These had enhanced gene expression ranging from 40 to 4500-fold. For ease of analysis, they were categorized into four groups: G1-G4, based on relative expression levels, with G1 (<500-fold), G2 (>500, <1000-fold), G3 (>1000, <2000-fold), and G4 (>2000-fold). 024_OE plants had thinner and weaker aerial parts with delayed growth (**Supplemental Figure 15B-J**). Severity of this phenotype intensified with higher expression levels. The plants had reduced height, stem width, leaf width, and root length. The leaves of *ONAC024* overexpressing plants displayed a drooping nature due to a decrease in the number of minor veins (**Supplemental Figure 16A-E**), while the number of major veins remained unchanged. Flowering was significantly delayed or absent in the plants overexpressing *ONAC024*, except for the OE_024_1 line, which had comparatively lower expression levels. Hence, ectopic overexpression of ONAC024 was detrimental to plant growth.

*ONAC024* and its alternatively spliced isoforms, were further functionally characterized (**Figure 3A**). *ONAC024* was overexpressed in a seed-preferential manner (024_SOE). *ONAC024* knock-down transgenic plants (024_KD) were raised using miRNA mediated gene silencing and RNAi mediated gene silencing (Reynolds et al., 2004; Warthmann et al., 2008) (**Supplemental Figure 17**). To identify the differential role of *ONAC024* alternatively spliced isoforms in regulating rice grain size, CRES-T technique was applied (Mitsuda et al., 2011), where each of ONAC024_A, ONAC024_B and ONAC024_C were converted into a chimeric repressor by fusion with EAR motif. The CRES-T lines were named as 024_A_CRES-T, 024_B_CRES-T and 024_C_CRES-T, respectively. The transactivation assay of CRES-T constructs in yeast (**Supplemental Figure 18)** showed that the constructs had minimal activation ability unlike ONAC024. Two independent lines from each transgenic plant type were analysed for the functional characterization of *ONAC024*. The S3 grains (stage with maximum *ONAC024* expression) of 024_SOE showed up to 6-10 fold upregulation and 024_KD plants showed 4-10 fold downregulation of ONAC024 (**Supplemental Figure 19).**

**Figure 3.**
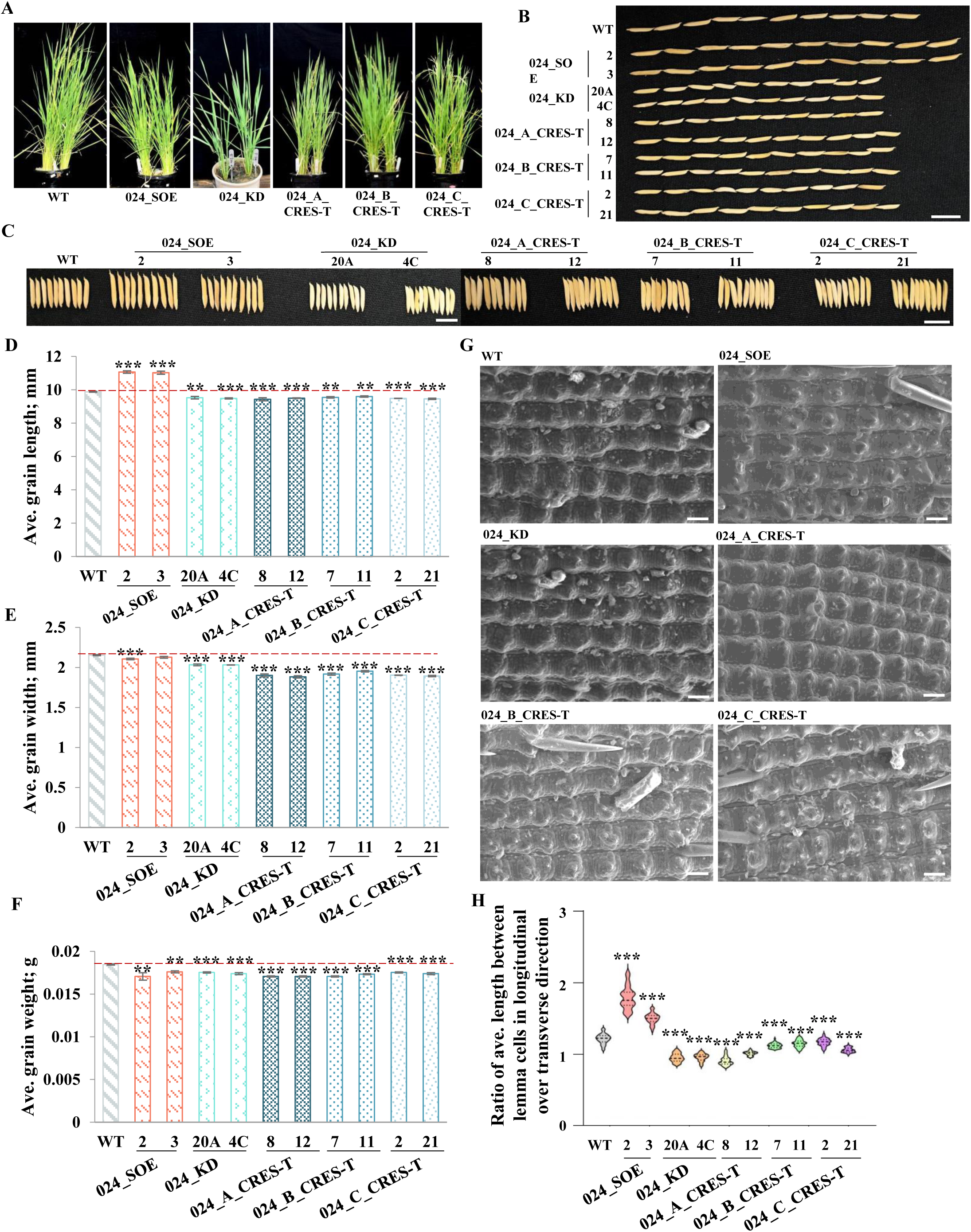
*ONAC024* has a positive effect on grain length. **(A)** *ONAC024* seed-preferential overexpression (024_SOE), knock-down (024_KD) and CRES-T generated (024_A_CRES-T, 024_B_CRES-T and 024_C_CRES-T for all three alternatively spliced forms of *ONAC024*) transgenic rice plants in comparison with wild type (WT) control. **(B)** and **(C)** Mature grains from WT and two independent lines each from different *ONAC024* transgenic plants (024_SOE – T3 grains; 024_KD – T4 grains; 024_A/B/C_CREST – T3 grains) representing **(B)** grain length and **(C)** grain width phenotypes, respectively. Scale bar, 1 cm. **(D)** to **(F)** Bar graphs show variation in **(D)** average per grain length, **(E)** average per grain width and **(F)** average per grain weight between WT and *ONAC024* transgenic grains (n = 50*3). The data was calculated from grains of three independent plants for each transgenic line. **(G)** SEM images from the upper surface of lemma to compare seed husk morphology between WT and ONAC024 transgenic grains. Scale bar, 20 µm. **(H)** Violin plot shows ratio of average length between lemma cells in longitudinal over transverse direction in 024_SOE, 024_KD, 024_A_CRES-T, 024_B_CRES-T and 024_C_CRES-T grains as compared with WT, n=45 (15*3). Student’s t-test was used for calculating the significance (**\***p ≤ 0.05; **\*\***p ≤ 0.01, **\*\*\***p ≤ 0.005).

In terms of grain size traits, the length of 024_SOE grains (**Figure 3B, D**) showed a significant increase (∼11.72%). Grain length was decreased in 024_KD (∼3.85%), 024_A_CRES-T (∼4.26%), 024_B_CRES-T (∼3.14%) and 024_C_CRES-T (∼4.16%) grains. SEM observations (**Figure 3G**; **Supplemental Figure 20A**) showed that the number of cells in the longitudinal direction were increased in all ONAC024 transgenic grains. However, a significant increase in average lemma cell length **(Supplemental Figure 20B)** in 024_SOE and decrease in the rest was observed. Hence, 024_SOE caused an increase in grain length by increasing both cell size and proliferation. A significant decrease in grain width (**Figure 3C, E**) was observed for 024_KD (∼5.76%), 024_B_CRES-T (∼12.26%), 024_B_CRES-T (∼9.48%) and 024_C_CRES-T (∼12.26%) when compared with WT. The number of cells in the transverse direction was increased in 024_KD, 024_A_CRES-T, 024_B_CRES-T and 024_C_CRES-T (**Supplemental Figure 20C)**. The average lemma cell width (**Supplemental Figure 20D)** was decreased only in 024_B_CRES-T and 024_C_CRES-T grains, accounting for the decreased width. The ratio of average length between lemma cells in the longitudinal direction over transverse direction (**Figure 3H**) was increased in 024_SOE, implying wider husk seeds; and decreased in the rest. Grain weight (**Figure 3F**) was significantly decreased in all lines, 024_SOE (∼6.17%), 024_KD (∼5.417%), 024_A_CRES-T (∼7.58%), 024_B_CRES-T (∼6.82%) and 024_C_CRES-T (∼5.74%) as compared to WT grains. These results suggest that *ONAC024* is a positive regulator of grain length, and variants ONAC024_B and ONAC024_C positively regulate (since these are CRES-T constructs) both grain length and width by regulating both cell size and cell number. Hence, ONAC024 and ZOS1-15 exert an opposite grain phenotype.

The height, number of tillers and panicles per plant were significantly decreased of all transgenic plants for ONAC024 (**Figure 3A**, **Supplemental Figure 21A, B, C)**. Flag leaf length was increased significantly only in 024_B_CRES_T plants (**Supplemental Figure 21D**). Panicle architecture was also affected (**Supplemental Figure 22A-D)**. The panicle length was decreased significantly in 024_SOE, 024_KD and 024_B_CRES_T plants **(Supplemental Figure 22B).** Number of primary branches was significantly decreased in 024_SOE, 024_KD, 024_A_CRES_T and 024_B_CRES_T plants (**Supplemental Figure 22C**) but the number of secondary branches was significantly decreased only in 024_C_CRES_T plants **(Supplemental Figure 22D).** The number of filled grains per plant was significantly decreased by 74.98% - 77.80% in all these transgenic plants (**Supplemental Figure 22E**), resulting in decreased yield per plant (**Supplemental Figure 22G**). However, the number of unfilled grains per plant was significantly increased only in 024_SOE plants **(Supplemental Figure 22F).**

### *ZOS1-15* and *ONAC024* regulate starch accumulation

Mature grains of 1-15_KD, 1-15_KO and 024_SOE had a translucent endosperm similar to WT (**Figure 4A**; **Supplemental Figure 23**). Concomitantly, the apparent amylose content (AAC) (**Figure 4B**) was increased in these grains by 22%, 26.7% and 27.8%, respectively, compared to WT. The dark blue color in the colorimetric assay (**Figure 4C**) for these grains implied increased amylose content. The central endosperm (**Figure 4D**) in these three grains consisted of more densely packed, polyhedral starch granules similar to those of WT endosperm. On the other hand, 1-15_OE, 024_KD, 024_A_CRES-T, 024_B_CRES-T and 024_C_CRES-T grains exhibited a visible increase in chalkiness when compared with WT (**Figure 4A**; **Supplemental Figure 23**). The AAC (**Figure 4B**) was decreased by 40.6%, 45.5% and 41.1% in 1-15_OE, 024_KD, 024_A_CRES-T grains, respectively. It was unaffected in 024_B_CRES-T and 024_C_CRES-T grains. The AAC was also confirmed by iodine colorimetric method (**Figure 4C**) where a lighter blue color in 1-15_OE, 024_KD and 024_A_CRES-T indicated decreased AAC. Under SEM (**Figure 4D**), the transverse sections from the central region of grains of 1-15_OE, 024_KD, 024_A_CRES-T, 024_B_CRES-T and 024_C_CRES-T grains was seen to be filled with more loosely packed, irregularly shaped spherical starch granules corresponding to the chalky region of the endosperm. In terms of expression of starch related genes (**Figure 4E**; **Supplemental Figure 24)**, the expression of *GPT* was significantly downregulated in both 1-15_OE and 024_SOE grains. Few genes such as *MST4, AGPS1, AGPL1* and *SSIIIa* had similar regulation upon increased expression of *ZOS1-15* or decreased expression of *ONAC024*, and behaved inversely upon their opposite expression. Amongst the alternatively spliced isoforms of ONAC024, the CRES-T plants of the main form ONAC024_A behaved like the 024_KD plants with lower amylose content. However, ONAC024_B and ONAC024_C did not affect the amylose content. However, the pattern of chalkiness was different amongst the three variants of ONAC024 and this can also be justified by the differential regulation of starch biosynthesis genes in the grains of CRES-T lines.

**Figure 4.**
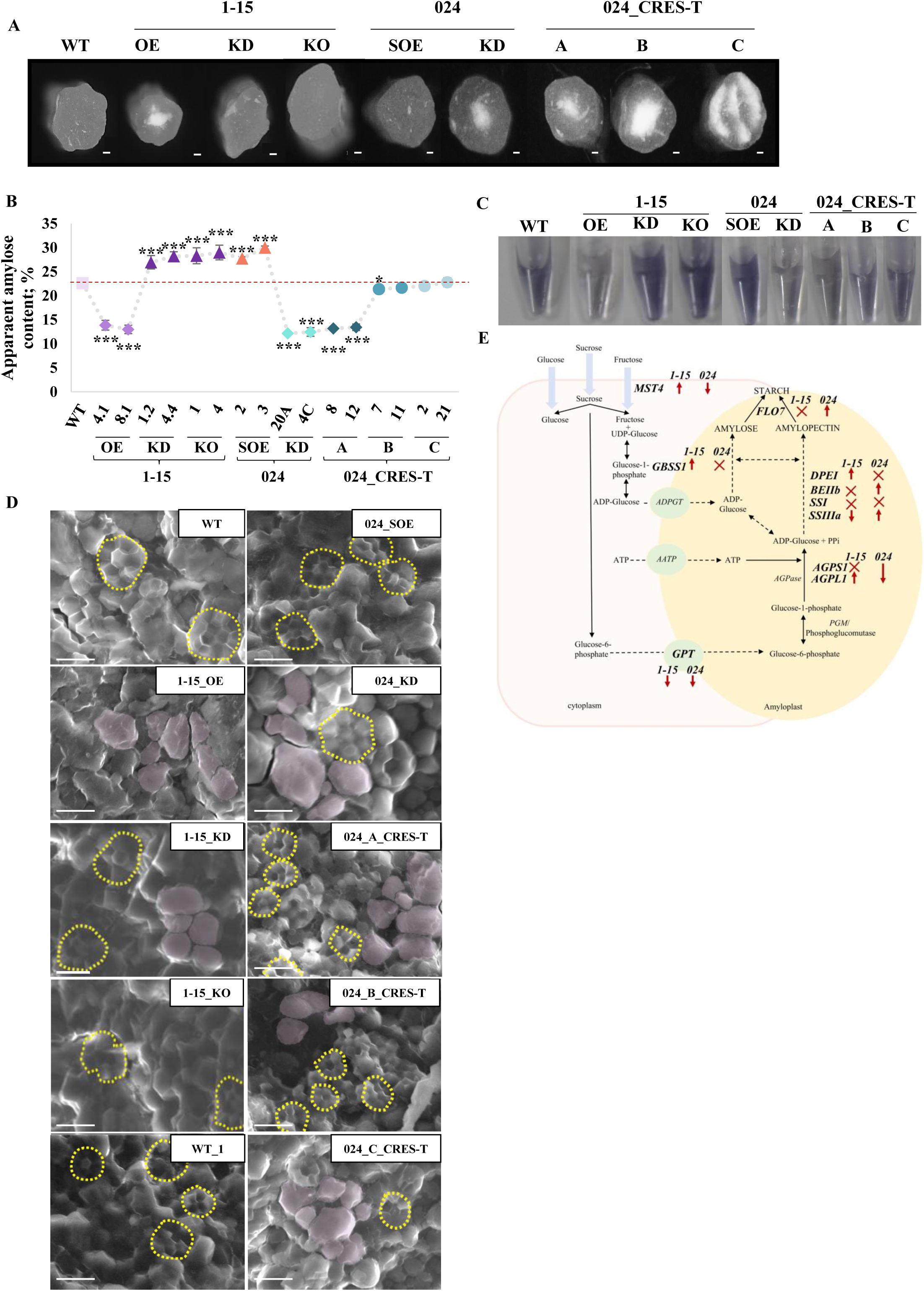
Knock-down/ knock-out of *ZOS1-15* and seed-preferential overexpression of *ONAC024* have a positive influence over amylose. **(A)** Hand sections of WT and transgenic grains of *ZOS1-15* and *ONAC024* to show grain chalkiness. Scale bars, 1 mm. **(B)** Line graph shows AAC in WT, *ZOS1-15* and *ONAC024* transgenic and mutant grains. The data was calculated from three biological replicates each from two independent transgenic lines of *ZOS1-15* and *ONAC024* (Student’s t-test at *p ≤ 0.05; ***p ≤ 0.005). **(C)** Iodine colorimetric assay to determine amylose content in grains of WT, ZOS1-15 and ONAC024. **(D)** Morphology of starch granules in grains of *ZOS1-15* and *ONAC024* along with WT. The polyhedral and irregularly shaped starch granules have been shown in yellow (dotted line) and pink color, respectively. Scale bars, 10 µm. **(E)** Representation of the starch biosynthesis pathway to show the differential expression of participating genes in 1-15_OE and 024_SOE grains. The up and down arrows indicate upregulation and downregulation, respectively. Cross indicates no significant variation in expression, with respect to WT.

Monosaccharide transporter, *MST4* expression was downregulated in 1-15_KD, 1-15_KO and 024_SOE grains and upregulated in 1-15_OE grains. The expression of *AGPS1* and *AGPL1* was significantly downregulated in 024_SOE grains, and upregulated in 1-15_KD and 1-15_KO. The gene encoding for starch branching enzyme (*BEIIb*) was upregulated only in 024_SOE grains. Amylopectin synthesis-related gene *SSIIIa* transcript level was decreased in 1-15_KO and increased in 1-15_KD and 024_SOE grains. The expression of *SSI* and amylose synthesis gene, granule bound starch synthase (*GBSSI*) was upregulated in 1-15_KO and remained unaffected in 1-15_KD and 024_SOE grains. *FLO7* expression was increased in 1-15_KO, 024_SOE and decreased in 1-15_KD grains. These results highlighted the role of *ZOS1-15* and *ONAC024* in the regulation of starch biosynthesis in rice grains and imply that ZOS1*-*15 negatively influences amylose production while ONAC024 increases it. This data further emphasizes on the role of the repressor-activator module of these TFs functional in rice grain development.

### Total protein content is positively regulated by ONAC024 and negatively by ZOS1-15

In terms of protein accumulation in *ZOS1-15* and *ONAC024* transgenic seeds relative to WT, a significant decrease in total protein content (TPC) was observed in 1-15_OE (∼34%) and 024_C_CRES-T (∼6.55%) seeds while 1-15_KD (∼14.36%), 1-15_KO (∼13.68%), 024_SOE (∼12.84%), 024_KD (∼5.94%) and 024_A_CRES-T (∼5.61%) had a higher TPC. It remained unaffected in 024_B_CRES-T seeds (**Figure 5A**). Rice contains 7% to 10% TPC of the grain dry weight (Shende et al., 2019). Out of this, glutelins (80%), prolamins (20-25%), albumins (4– 22%) and globulins (5–13%) account for major seed storage proteins (SSPs) (Hoogenkamp et al., 2017). On the SDS-PAGE (**Figure 5B-D**), 13 kDa prolamin bands were thicker from 1-15_KD, 1-15_KO, 024_SOE and 024_A_CRES-T grains. This correlated with high expression of *PRO16* and *PRO18* in these seeds (**Figure 5E**). In addition, 024_C_CRES-T and 024_KD showed low intensity bands for 13 kDa prolamins (**Figure 5C, D**). 024_SOE showed a decreased intensity of glutelin precursor (∼55 kDa) but additional bands for glutelin acidic (∼35 kDa) and basic subunits (∼25 kDa) that correlates with high expression of glutelins (*GLU6*, *GLU14* and *GLU19*) (**Figure 5C, E**). On the other hand, 024_KD showed a decrease in the band intensity of glutelin acidic (∼35 kDa) and basic subunits (∼25 kDa) that corresponded to decreased expression of glutelins (*GLU6*, *GLU14* and *GLU19*) (**Figure 5C, E**). 024_A_CRES-T showed high intensity bands for rice allergen proteins that coincided with an increased expression of globulins (*GLB1, GLB2*) (**Figure 5D, E**). 024_SOE and 1-15_OE showed opposite SSP expression patterns while 1-15_OE and 024_KD showed similar patterns of expression for *ALB3*, *ALB13*, *ALB16, GLB1, GLB2, GLB4, GLU6, GLU14, GLU19, PRO16, PRO18*. 1-15_OE and 1-15_KD showed opposite expression pattern for *ALB13, ALB16, GLB1*. Opposite expression pattern for *GLB2, GLB4, GLU14, GLU19, PRO16* was observed in 1-15_KD and 024_SOE. 024_KD and 1-15_KO showed opposite expression pattern for *GLB4, GLU6, GLU19, PRO16* and *PRO18*. 1-15_KO and 024_SOE showed similar expression pattern for *GLB4*, *GLU6*, *GLU19*, *PRO16* and *PRO18*. These results imply that amongst the main components of the repressor-activator module, ZOS1-15 is a negative regulator of TPC while, ONAC024 is a positive regulator. Amongst the spliced variants, ONAC024_B does not affect TPC, while ONAC024_C is also a positive regulator of TPC.

**Figure 5.**
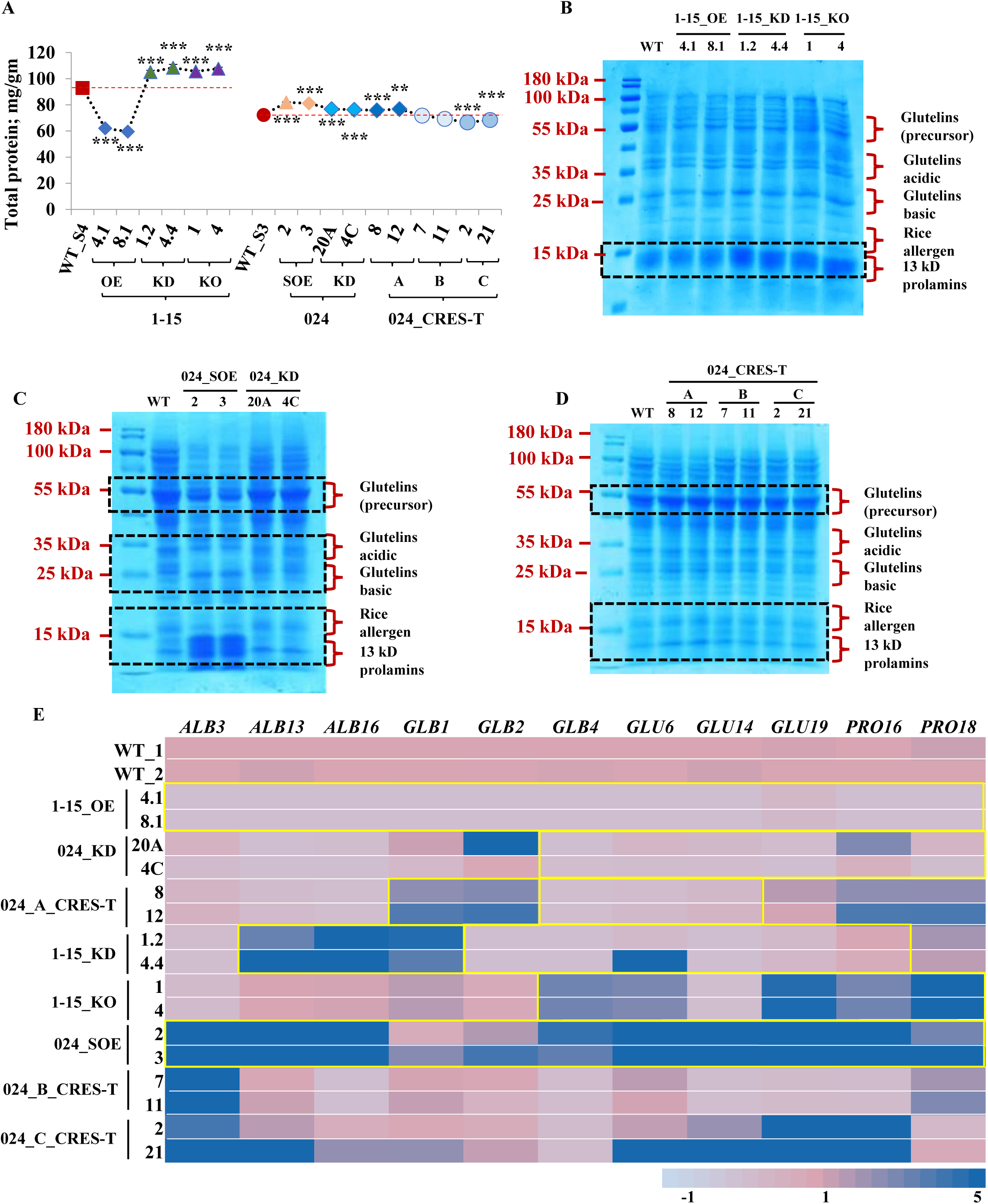
*ZOS1-15* and *ONAC024* control grain total protein content in an opposite manner. **(A)** TPC estimation from seeds of *ZOS1-15* (S4 stage), *ONAC024* (S3 stage) transgenic, mutant and WT plants. WT_S4 and WT_S3 indicate WT grains for each stage. Asterisks denote significant difference as determined by Student’s t-test (**\*\***,**\*\*\*** = p value ≤ 0.01, ≤ 0.005 respectively). Error bars = ± SE. SDS-PAGE of total protein extracted from **(B)** seeds of WT and *ZOS1-15* plants and **(C-D)** seeds of WT and *ONAC024* transgenic plants. SDS-PAGE gels show prominent bands for glutelin precursor (∼50 kDa), glutelin acidic subunits (∼30 kDa), glutelin basic subunit (∼22 kDa), rice allergen (14-16 kDa) and 13 kDa prolamins which are indicated on the right side of the gel. Black boxes highlight bands with intensity variation between WT and different types of transgenic plants. **(E)** Heat map shows differential expression of *SSP* genes in *ZOS1-15* and *ONAC024* grains relative to the expression levels in WT_1 and WT_2 respectively, analyzed by qRT-PCR. Yellow outline indicates an opposite expression pattern of the gene between *ZOS1-15* and *ONAC024* grains. Color bar at the bottom of the heat map represents fold change, thereby left and right values of the color bar indicate low and high expression, respectively.

### ZOS1-15 and ONAC024 regulate expression of the same downstream genes

Out of all the SSPs whose expression was tested, two genes *GLU6* (**Supplemental Figure 25A**) and *ALB13* (**Supplemental Figure 25B)** were taken for further analysis. These were significantly downregulated in 1-15_OE, 024_KD, 024_A_CRES-T seeds and upregulated in 1-115_KD, 1-15_KO, 024_SOE seeds. Therefore, to understand the regulation of these genes by ZOS1-15 and ONAC024, a yeast one-hybrid assay was performed with *ProGLU6* (**Supplemental Figure 25C**) and *ProALB13* (**Supplemental Figure 25D**). The expression of *Aur^R^*leading to growth of yeast cells on the SD media lacking leucine, uracil and supplemented with 200 ng/mL Aureobasidin demonstrated the binding of ZOS1-15 and ONAC024 to *ProGLU6* and *ProALB13*. Yeast one-hybrid assays with the deletion constructs of these promoters showed that the binding occurred at *ProGLU6_1* (-1771 bp) and *ProALB13_3* (-974 bp) for ZOS1-15 and *ProGLU6_4* (-606 bp) and *ProALB13_3* (-830 bp) for ONAC024. ONAC024_B and ONAC024_C isoforms did not bind to *ProGLU6* (**Supplemental Figure 25C**) and *ProALB13* **(Supplemental Figure 25D)**. The NAC domain of ONAC024 was found to be responsible for binding to *ProGLU6* at *ProGLU6_4* (**Supplemental Figure 25C**) and *ProALB13* at *ProALB13_3* (**Supplemental Figure 25D)**. The sequence of zinc finger binding site (ZF_BS) was same for both *ProGLU6* and *ProALB13* (**Figure 6A, B**) while NAC binding site (NBS) had different sequences in both promoters. To validate the binding sites, EMSA was performed using biotin labelled 2X ZF_BS for *ProGLU6* and *ProALB13* and 2X/3X_NBS for *ProGLU6* and *ProALB13*, respectively, as the probe, and purified GST_ZOS1-15 and MBP_ONAC024. A clear shift in EMSA confirmed the binding of ZOS1-15 and ONAC024 to *ProGLU6* (**Figure 6A, B**) and *ProALB13* (**Figure 6C, D**), respectively. The binding was reduced with increasing concentrations of unlabeled probe. No shift was detected with mutated probes of *ProGLU6* and *ProALB13*. These results confirmed that ZOS1-15 and ONAC024 specifically bound to the predicted ZF_BS and NBS on *ProGLU6* and *ProALB13*.

**Figure 6.**
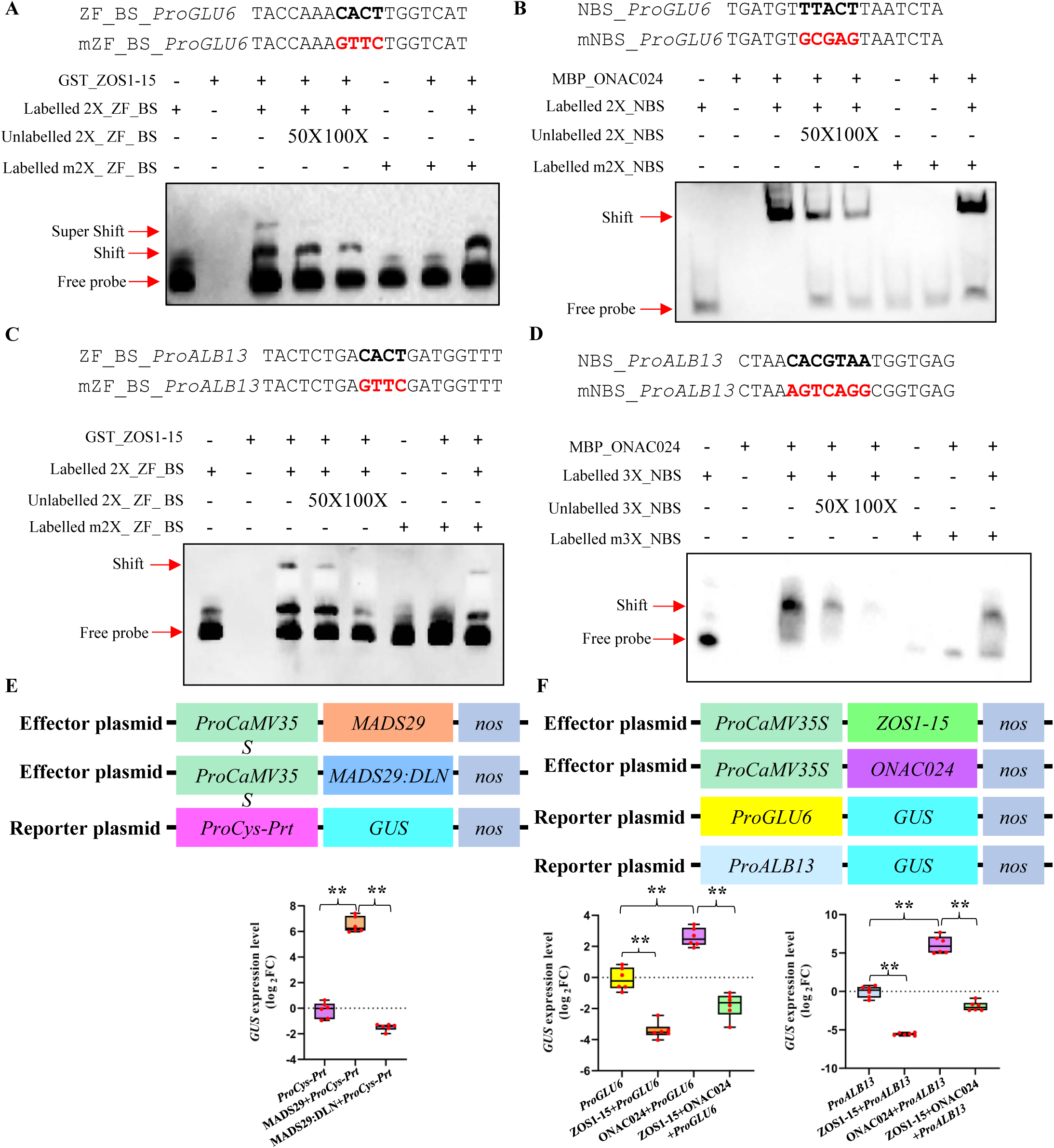
ZOS1-15 and ONAC024 bind to the promoters of and regulate *SSP* expression in an antagonistic manner. **(A)** to **(D)** EMSAs show the binding of ZOS1-15 and ONAC024 to the biotin labelled C_2_H_2_ zinc finger binding site (ZF_BS) and NAC binding site (NBS), respectively, present in the promoters of **(A, B)** *GLU6* and **(C, D)** *ALB13*. The sequences of 1X ZF_BS and NBS have been highlighted in black and the corresponding mutated region is in red. The efficiency of binding was checked with two different concentrations of unlabeled probe 50X and 100X. **(E)** and **(F)** Reporter effector assays showing *in planta* validation of **(E)** DLN motif from ZOS1-15 as a transcriptional repressor and **(F)** suppression of transcriptional activation of ONAC024 by ZOS1-15 on the promoter of *GLU6* and *ALB13*. Top panel is the schematic representation of constructs used. The effector plasmids have **(E)** MADS29 or MADS29 fused with DLN motif from ZOS1-15 (MADS29:DLN) and **(F)** ZOS1-15 or ONAC024, under the control of *CaMV35S* promoter. The reporter plasmid has *β-glucuronidase* (GUS) gene driven by the promoter of **(E)** *Cys-Prt* and **(F)** *GLU6* or *ALB13.* Median levels were calculated from six biological replicates with statistical significance using Student’s t-test at **\*\***p ≤ 0.01.

ZOS1-15 and its DLN repressor motif (DLNYPP) have been shown to strongly repress reporter gene activity in yeast-based assays (Singh et al., 2019). *In planta* reporter effector assay showed that this DLN motif could decrease the activation ability of MADS29 (**Figure 6E**) on *Cys-Prt* promoter to which it has been shown to bind (Yin and Xue, 2012). It has been shown earlier that a mutated version of DLNYPP does not cause repression in yeast (Singh et al., 2019). This confirms that the DLN motif of ZOS1-15 imparts it the repressive ability. Another set of reporter effector assays was done to test the transcriptional activity of ZOS1-15 and ONAC024 on the promoters of *GLU6* and *ALB13.* When the effector was ZOS1-15, the expression of GUS driven by either *ProGLU6* or *ProALB13* (**Figure 6F**) was highly downregulated. This also confirmed that ZOS1-15 acted as a repressor on these promoters. As an effector, ONAC024 could strongly activate *GUS* expression (**Figure 6F**) from *ProGLU6* and *ProALB13*. Next, the role of ZOS1-15_ONAC024 dimer in the regulation of *SSP* promoter was elucidated by *in planta* reporter effector assay. The high transcriptional activity of ONAC024 on both the promoters was significantly decreased in the presence of ZOS1-15. This implied that ZOS1-15 as a repressor could leverage the activation of *SSP* promoters by ONAC024.

### Other NAC TFs form a part of the repressor-activator module

Interactions amongst various NAC TFs have already been reported in different plant systems including rice (Mendes et al., 2013; Mathew et al., 2016). Interaction between ONAC024 and ONAC023 was tested through yeast two-hybrid assay, *in planta* BiFC **(Supplemental Figure 26A, B, C)**, pull down and *in planta* CoIP assays **(Figure 7A, B)**. Growth of yeast colonies on the SD/-AHLT plate and the presence of blue colour on SD/-AHLT + X-α-gal showed their interaction. The presence of YFP signal in the *Nicotiana* leaves by BiFC assay indicated that ONAC024 physically interacts with ONAC023 and the dimer is localized in the nucleus. The TRR domain of ONAC024 was responsible for this interaction. ONAC024_C and ONAC024_B showed decreasing levels of interactions, respectively, which was also seen in the yeast two-hybrid assay with their TRR domains. *In vitro* pull-down assay was performed using bacterial expressed proteins, MBP_ONAC024 and GST_ONAC023. Immunoblotting with anti-MBP and anti-GST antibodies, separately, detected the presence of MBP_ONAC024 (76.91 kDa) and GST_ONAC023 (53.97 kDa), respectively. The interaction was also validated through *in-planta* CoIP assay by the detection of YFP_ONAC024 (62.54 kDa) (**Figure 7B**).

**Figure 7.**
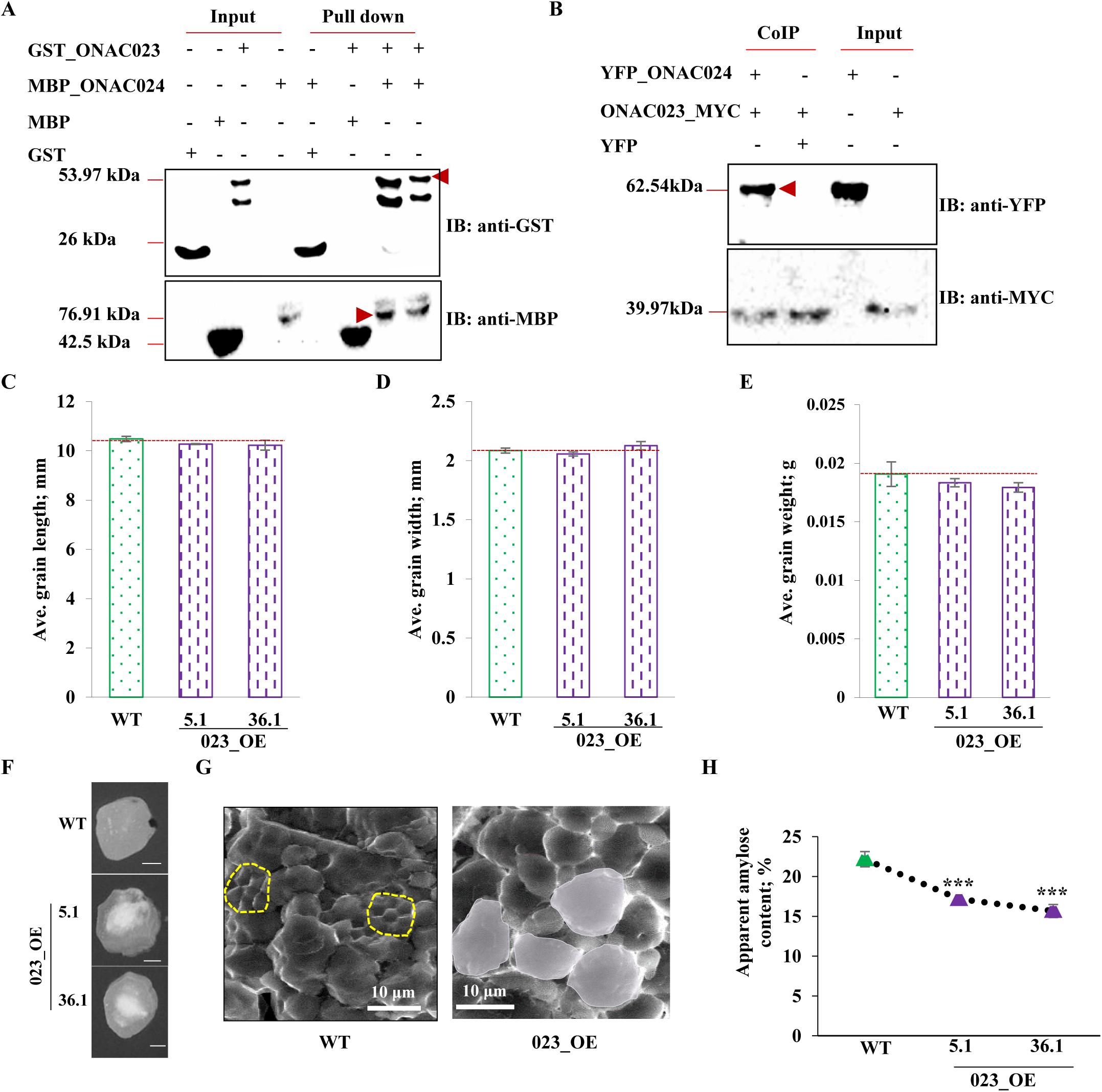
ONAC023 interacts with ONAC024 and negatively regulates amylose content. **(A)** Pull down assay showing interaction of MBP_ONAC024 (76.91 kDa) with GST_ONAC023 (53.97 kDa). **(B)** *In-planta* co-immunoprecipitation assay confirming the interaction of YFP_ONAC024 (62.54kDa) with ONAC023_MYC (39.97kDa). Pull down and Co-IP lanes with expected protein bands have been indicated by red arrows. IB denotes immunoblot. Graphs representing reproductive traits in 023_OE lines compared to WT; **(C)** average per grain length; **(D)** average per grain width and **(E)** average per grain weight (n = 50*3). **(F)** Transverse sections representing grain chalkiness in wild type (top), and 023_OE grains (5.1 – T3 grains; 36.1 – T4 grains); scale bar = 100µm. **(G)** Structural analysis of starch granules in WT and 023_OE lines by SEM, scale bar = 10µM; polyhedral starch granules have been marked in yellow and irregular shaped ones are in pink. **(H)** Estimation of AAC in mature seeds of WT and ONAC023 OE lines. Student’s t-test was used for calculating the significance (**\*\*\*** = p-value< 0.005).

qRT-PCR analysis of *ONAC023* in various vegetative and reproductive tissues showed its highest expression in seed developmental stages from S2 to S5 **(Supplemental Figure 27A)**. To investigate the function of *ONAC023*, overexpression transgenic lines were generated (023_OE) **(Supplemental Figure 27B).** qRT-PCR confirmed overexpression of *ONAC023* in these lines **(Supplemental Figure 27C).** 023_OE plants had decreased height, flag leaf length and width **(Supplemental Figure 27D-F).** They flowered late by 2-3 days as compared to WT plants **(Supplemental Figure 27G)**. Panicle length, number of primary and secondary branches in the panicle were decreased **(Supplemental Figure 27H-J, N).** Number of filled grains and the yield was decreased and number of unfilled grains was increased **(Supplemental Figure 27K-M).** However, no significant change was observed for grain length, grain width and grain weight in 023_OE plants (**Figure 7C-E**; **Supplemental Figure 27O, P)**. Transverse sectioning of 023_OE mature grains exhibited higher endosperm chalkiness in comparison to wild type (**Figure 7F**). SEM imaging of 023_OE seeds confirmed the presence of irregular starch granules unlike the polygonal shaped granules observed in WT seeds (**Figure 7G**). AAC was reduced by 22-31% in these grains (**Figure 7H**, **Supplemental Figure 27Q).** However, TPC showed no change in these grains **(Supplemental Figure 28A, B)**. In our previous study, using yeast two-hybrid assay and BiFC, we have reported that ONAC023 interacts with ONAC026, which also interacts with ONAC020. Each of these dimers are nuclear localized (Mathew et al., 2016). To establish other components of our repressor-activator module, we validated the above interactions through pull-down assay. Detection of GST_ONAC026 (63.4 kDa) in the MBP_ONAC023 bound fraction and MBP_ONAC023 (71.3kDa) in the GST_ONAC026 bound fraction confirmed the interactions **(Supplemental Figure 29A)**. Similarly, the interaction of ONAC026 with ONAC020 was also confirmed **(Supplemental Figure 29B)**. Amongst the module components determined till now, no interaction was observed between ZOS1-15 and ONAC023; ONAC024 and ONAC20; ONAC24 and ONAC26; ZOS1-15 and ONAC20; ZOS1-15 and ONAC026 and ONAC20 and ONAC023 **(Supplemental Figure 30A, B)**. Hence, ONAC024, ONAC023, ONAC026 and ONAC020 interact in tandem with each other to form a functionally active complex to regulate various aspects of rice seed. In all probability, the ONAC024-ONAC023 association influences AAC.

### SS1/ ONAC025 competes with ONAC024 in regulating the synthesis of *SSPs*

Yeast one-hybrid assay identified the binding of another NAC TF, SS1/ ONAC025 to the 2 kb promoter region as well as the deletion fragments of *GLU6* and ALB13 at *ProGLU6_4* (-606 bp) (**Supplemental Figure 31A)** and *ProALB13_3* (-830 bp) (**Supplemental Figure 31B**). Further, the shift in EMSA, performed between SS1/ ONAC025 and NBS, confirmed its binding to the NBS of *ProGLU6* (**Figure 8A**) and *ProALB13* (**Figure 8B**). Further, *in planta* reporter effector assay showed that SS1/ ONAC025 suppressed the expression of *GUS* reporter driven by *ProGLU6* (**Supplemental Figure 32A**) and *ProALB13* (**Supplemental Figure 32B**). Due to the same binding site of ONAC024 and SS1/ ONAC025 for *ProGLU6* and *ProALB13*, competition EMSA was performed to test the influence of the presence of ONAC024 and SS1/ ONAC025 on each other in regulating these *SSPs* expression (**Figure 8C, D**). When equal amounts of both proteins were present, the relative band density for *ProGLU6* (**Supplemental Figure 33A**) or *ProALB13* (**Supplemental Figure 33B)** probes was comparable to that for a single protein, indicating competition between the two NACs. As the concentration of one protein was increased to 50X, the binding intensity increased. The relative band density was highest when 100X concentration of SS1/ ONAC025 was incubated with *ProGLU6* (**Supplemental Figure 33A**) or *ProALB13* (**Supplemental Figure 33B)** probes along with ONAC024. The band intensity was lesser when 50X or 100X concentrations of ONAC024 were incubated with *ProGLU6* or *ProALB13* probe along with SS1/ ONAC025. This indicated a stronger binding of SS1/ ONAC025 onto the NBS. To substantiate the competition between ONAC024 and SS1/ ONAC025, *in planta* reporter effector assays were performed in *N. benthamiana* leaf epidermal cells with *GUS* reporter gene driven by target promoters, *ProGLU6* and *ProALB13* (**Figure 8E, F**). Individually, ONAC024 activated while SS1/ ONAC025 repressed *GUS* expression. When *ONAC024* and *SS1/ ONAC025* were co-expressed, SS1/ ONAC025 could repress the transcriptional activation by ONAC024 on *ProGLU6* and *ProALB13*. This indicates that SS1/ ONAC025 interferes with binding of ONAC024 to *ProGLU6* and *ProALB13* at the NBS.

**Figure 8.**
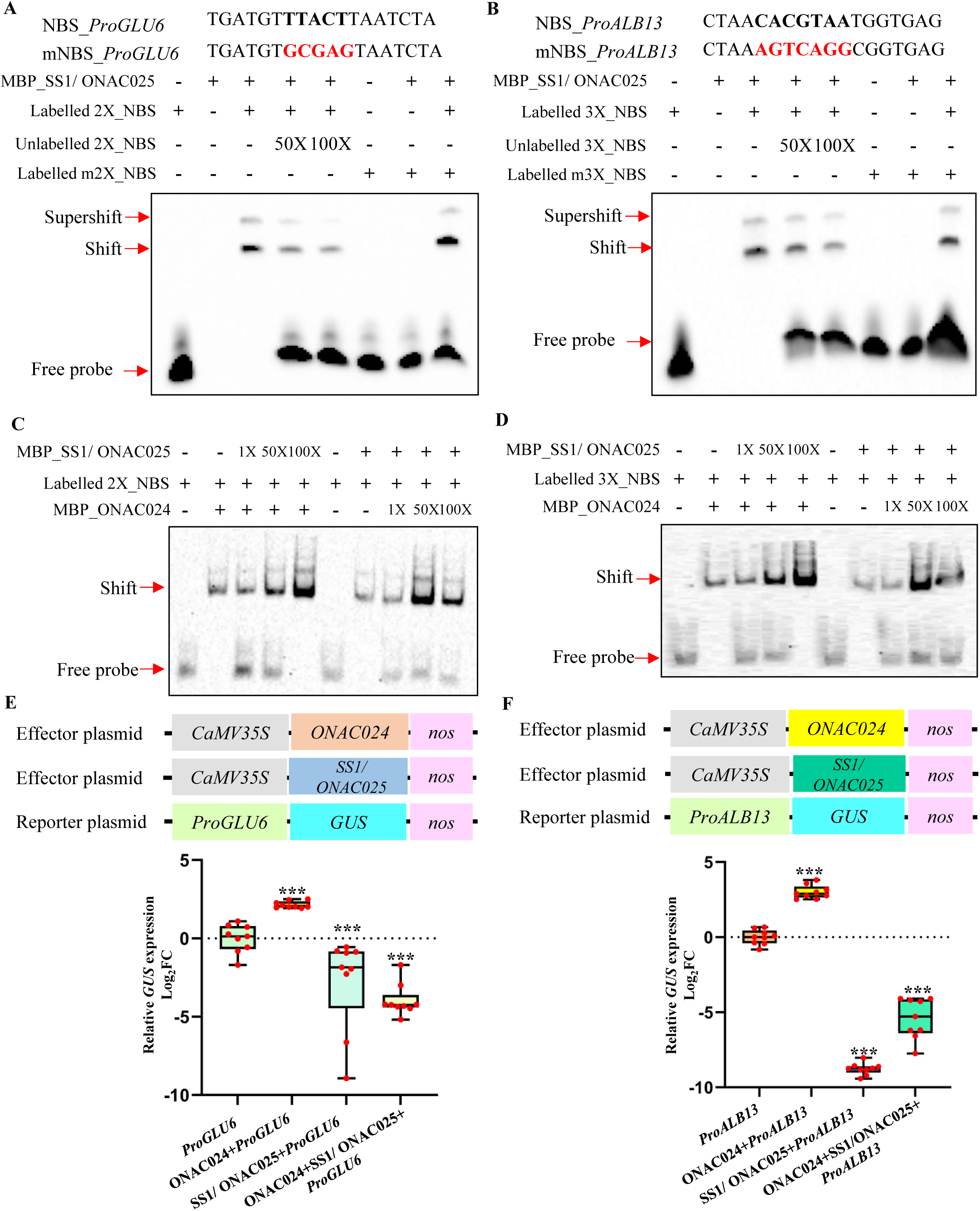
SS1/ ONAC025 competes with ONAC024 for the same NBS and regulates *SSP* expression in an opposite manner. **(A)** and **(B)** EMSAs demonstrating the direct binding of SS1/ ONAC025 onto the NBS of **(A)** *GLU6* and **(B)** *ALB13*. The sequence of NBS_*GLU6* or NBS_*ALB13* and its mutated probe, mNBS_Pro*GLU6* or mNBS_Pro*ALB13* are mentioned above the immunoblot, where black highlights NBS and red shows its mutated sequence. The binding was checked with two different concentrations of unlabelled probe 50X and 100X. The bound and free probes are marked. **(C)** and **(D)** EMSAs show the competition between SS1/ ONAC025 and ONAC024 onto the NBS present on the promoter of **(C)** *GLU6* and **(D)** *ALB13*. Labelled NBS probes with increasing concentrations (1X, 50X and 100 X) of either MBP_ONAC024 or MBP_SS1/ ONAC025 were incubated to confirm their competition onto the promoter of the two *SSPs*. The free probe and bound shift is highlighted on the left side of the blot. **(E)** and **(F)** Reporter effector assays to validate the role of SS1/ ONAC025 in interfering with the transcriptional activation ability of ONAC024 onto the promoters of **(E)** *GLU6* and **(F)** *ALB13*. The schematic representation of constructs used as reporter and effector plasmids in the assay has been shown above the dot-box plots. Dots signify biological replicates (n=9). One tailed Student’s t-test was used for calculating the significance at **\*\*\*** for p≤0.005.

### *SS1/ ONAC025* positively regulates grain yield, alters starch and protein accumulation

Given the competition of SS1/ ONAC025 with ONAC024, its function in rice was determined. Expression analysis of *SS1/ ONAC025* (**Supplemental Figure 34A)** showed its highest expression in S3 stage (∼700-fold). Our previous data suggests that though the ectopic expression of SS1/ ONAC025 results in lethal phenotype, it is a key regulator of rice grain filling and might control amylopectin production (Mathew et al., 2020). Therefore, to validate the role of *SS1/ ONAC025*, the gene was overexpressed in a seed-preferential manner and rice transgenic plants were raised and named as 025_SOE (**Supplemental Figure 34B)**. A significant overexpression (31-42-fold) (**Supplemental Figure 34C)** was observed in 025_SOE plants. Grain phenotyping of 025_SOE plants showed reduction in grain length by 3.14% **(Figure 9A, C)** and the SEM analysis of lemma cells of the spikelet hull **(Supplemental Figure 35A)** showed that both the number of cells in the longitudinal direction (15.8%) and the average lemma cell length (25.4%) were decreased (**Supplemental Figure 35B-C)**. Grain width was decreased by 9.76% (**Figure 9B, D**) but the number of cells in the transverse direction and the average lemma cell width (**Supplemental Figure 35D, E**) was unaffected in 025_SOE. Further, the ratio of average length between lemma cells in the longitudinal direction over transverse direction (**Supplemental Figure 35F**) was decreased by 12.1% in 025_SOE indicating less wider cells. The grain weight was also decreased (6.5%) (**Supplemental Figure 34K**). Hence, SS1/ ONAC025 negatively regulates grain size by affecting both cell size and proliferation. This is in contrast with the role of ONAC024 in grain size control.

**Figure 9.**
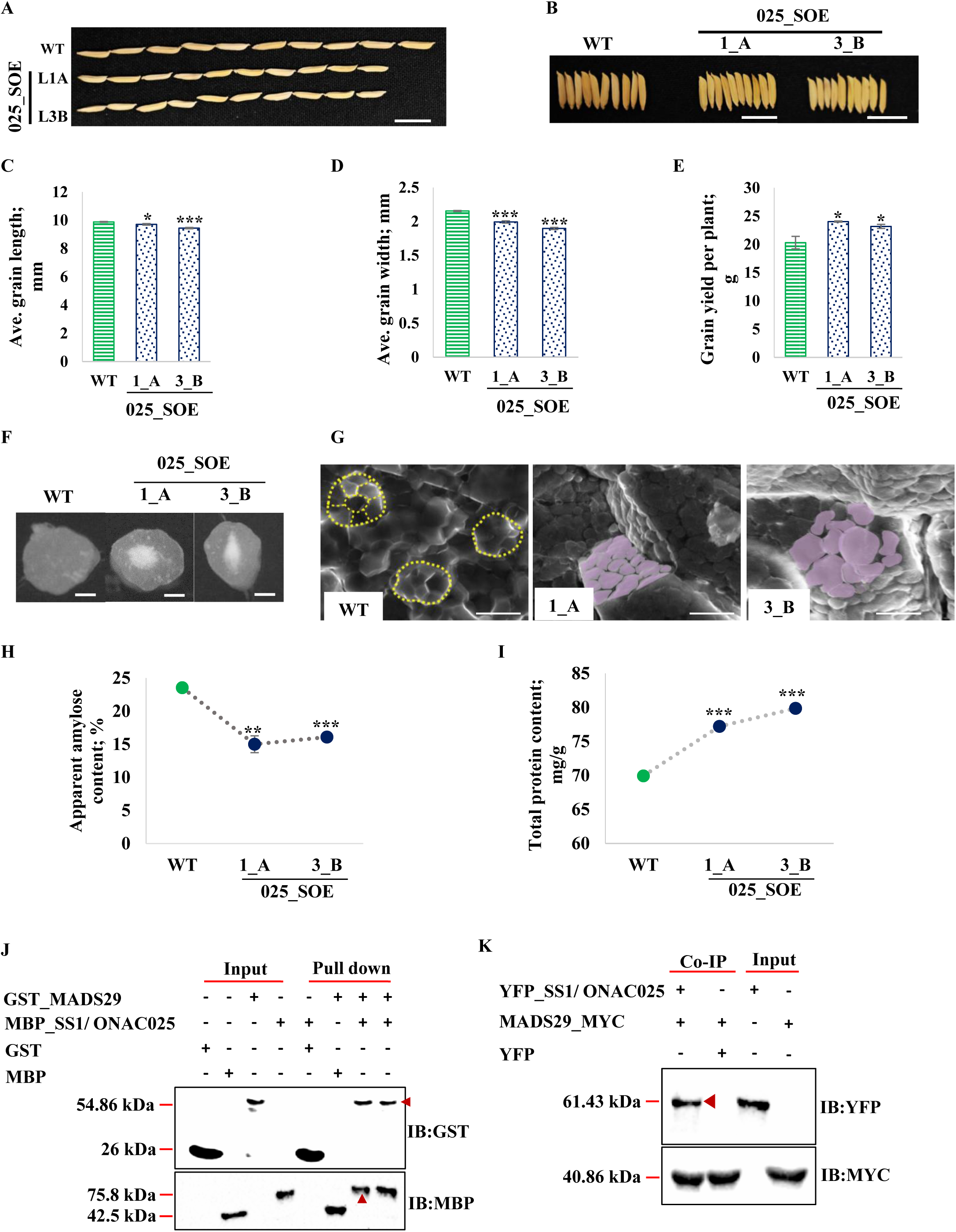
ONAC025 negatively regulates grain size and amylose but enhances yield and protein content; and forms a complex with MADS29. Mature grains of 025_SOE (T3 grains) and WT arranged **(A)** lengthwise and **(B)** widthwise. Bar, 1 cm. Bar graphs show **(C)** average per grain length, **(D)** average per grain width and **(E)** yield per plant in WT and 025_SOE plants, (n= 50*3). **(F)** Transverse sections of WT and 025_SOE grains under the stereomicroscope. Scale bars, 1 mm. **(G)** Morphology profile of compound-type starch granules in the central endosperm sections of 025_SOE mature grains in comparison with WT. The transverse hand-cut sections of grains have been viewed under SEM. The polyhedral starch granules are outlined with yellow dashed lines and irregular granules have been marked in pink. Scale bars, 10µm. Line graph shows **(H)** AAC in WT and SS1/ ONAC025 transgenic grains and **(I)** TPC of WT and 025_SOE grains. The data represents two independent transgenic lines with three biological replicates each. The graphs depict average ± SE and statistical significance by one-tailed Student’s t-test (**\***p<0.05, **\*\***p<0.01, **\*\*\***p<0.005). **(J)** Pull-down assay shows the interaction of MBP_SS1/ ONAC025 (75.8 kDa) with GST_MADS29 (54.86 kDa). The red arrow indicates the expected protein bands in the pull-down lane. IB, immunoblot. **(K)** Co-immunoprecipitation (Co-IP) assay confirming the interaction between SS1/ ONAC025 (61.43 kDa) and MADS29 (40.86 kDa). Input indicates total protein isolated from *N. benthamiana* leaves infiltrated with YFP_SS1/ ONAC025 and MADS29_MYC. The red arrow represents immunoprecipitated YFP_SS1/ ONAC025 in the protein mixture. IB denotes immunoblot.

Apart from grain size, vegetative and panicle traits were also affected. The plant height was decreased **(Supplemental 34B, D)** while the number of tillers and panicles (**Supplemental Figure 34E, G)** were increased in 025_SOE plants. Panicle architecture (**Supplemental Figure 34F**) in 025_SOE plants did not show significant variation in panicle length and number of secondary branches per panicle (**Supplemental Figure 34H, J**). However, the number of primary branches per panicle were decreased in 025_SOE plants as compared to WT (**Supplemental Figure 34F, I**). The total number of filled grains per plant was significantly increased by 18.1%-20.64% in 025_SOE plants (**Supplemental Figure 34L**), resulting in increased yield per plant (9.70%-12.81%) (**Figure 9E**). These results suggest that *SS1/ ONAC025* positively regulates tiller number and grain yield and negatively regulates grain size. These phenotypes are opposite to that observed for ONAC024_SOE plants.

Compared with WT, *025_SOE* grains exhibited increased chalkiness gathered towards the central endosperm (**Figure 9F**, **Supplemental Figure 36A**). AAC (**Figure 9H**, **Supplemental Figure 34M)** was significantly reduced (∼33.99%) in 025_SOE grains. SEM observations of transverse cut sections of 025_SOE (**Figure 9G**) revealed that the central endosperm was filled with irregularly shaped starch granules which corresponded with the chalky area. However, the starch granules in the peripheral region of endosperm (**Supplemental Figure 36B)** were densely packed and had a polyhedral shape in 025_SOE grains similar to WT, indicating the positional variations in starch granules within the endosperm. The expression of starch biosynthesis genes (**Supplemental Figure 36C)** was estimated. The expression of *GPT*, *AGPS1*, *AGPL3*, *SSI*, *SSIIIa* and *DPE1* were significantly upregulated in 025_SOE grains. In contrast, the expression of *MST4* and *GBSSI* were decreased. The differential expression of genes involved in starch synthesis indicates the involvement of *SS1/ ONAC025* in regulating amylose accumulation and starch structure in rice grain. TPC in 025_SOE grains (**Figure 9I**) was significantly increased (10.39-14.18%), a phenotype similar to 024_SOE. The total seed protein on 10% SDS-PAGE (**Supplemental Figure 37A)** indicated that the band intensity for glutelin acidic, glutelin basic, rice allergens and prolamins changed in 025_SOE grains. The expression of various SSPs (**Supplemental Figure 37B)** was quantified through qRT-PCR. The expression of *ALB3*, *ALB16, GLB1, GLB2, GLB3, GLB4* and *PRO18* was significantly upregulated. The expression of *ALB13, GLU6* and *GLU14* was decreased. *GLU19* and *PRO16* expressions were not altered in 025_SOE grains in comparison with WT. These results show that SS1/ ONAC025 regulates the total protein accumulation by altering the expressions of various *SSPs* in rice grain.

NAC TFs usually heterodimerize for stable DNA binding (Harrington et al., 2019). SS1/ ONAC025 however, did not show interaction with the other TFs, ZOS1-15, ONAC024, ONAC023, ONAC020 and ONAC026 (**Supplemental Figure 38**). Subsequently, SS1/ONAC025 interaction network was predicted by STRING database (Version 11.5) wherein MADS29 was highlighted. Yeast two-hybrid assay (**Supplemental Figure 39A**) showed that SS1/ ONAC025 interacted with MADS29. This interaction was confirmed through BiFC assay (**Supplemental Figure 39B)** where the reconstituted YFP signal was observed in the nucleus and cytoplasm. Further, the interaction was confirmed by *in-vitro* GST pull-down assay (**Figure 9J**) where the MBP_SS1/ ONAC025 (75.8 kDa) band in the pull-down lane when GST_MADS29 was bound; and the presence of GST_MADS29 (54.86 kDa) when MBP_SS1/ ONAC025 was bound, confirmed the interaction of SS1/ ONAC025 and MADS29. This interaction was also validated by *in planta* Co-IP assay (**Figure 9K**) where MADS29_MYC was able to immunoprecipitate YFP_SS1/ ONAC025 (61.43 kDa) and not the YFP control. These results indicated that MADS29 interacted with SS1/ ONAC025 and guided its nuclear localization. Summarily, SS1/ ONAC025 negatively regulates grain length, amylose content and positively regulates grain yield and TPC, and is a competitor to activator ONAC024 in regulating the synthesis of two *SSPs*.

## DISCUSSION

Being a staple crop, the rice grain has been a subject of intense research over generations. Many genes have been elucidated which regulate its size, the starch biosynthesis pathway has been determined and there is fair knowledge on seed storage proteins. However, information on genes which can connect all these aspects of the rice grain and their detailed molecular mechanisms has been lacking. The repressor-activator-competitor module reported in this paper consists TFs ZOS1-15-ONAC024-ONAC023-ONAC026-ONAC020 and SS1/ ONAC025 (**Figure 10**). In tandem or antagonistically, they regulate various aspects of rice development, especially grain size, AAC and TPC. This module consists of both positive and negative regulators of these traits. TFs often form a protein complex which consists of homo-or heteromers. This includes formation of a repressor-activator module, where a transcriptional repressor interacts with an activator to regulate the expression of downstream genes. Only a handful of repressor-activator interactions have been reported till date. Amongst these, a cascade involving DELLA-NAC regulates cellulose synthesis in rice (Huang et al., 2015). Repressor HvWRKY38 interacts with activator HvGAMYB and competes for binding to *α-amylase* promoter at different *cis*-elements (Zou et al., 2008). Repressor MaDof23 and activator MaERF9 interact and bind to different motifs, 5′-(T/A)AAAG-3′ and GCC box, respectively, on the promoters of ripening related genes to regulate their expression (Feng et al., 2016; Hao et al., 1998; Diaz et al., 2002). JA/ET-inducible activator OsERF87 and SA inducible OsWRKY76 repressor bind to promoter of *RSOsPR10* at GCC-*cis* element and W box, respectively, to maintain a balance between JA/ET and SA signaling in plants under stress conditions (Yamamoto et al., 2018).

**Figure 10.**
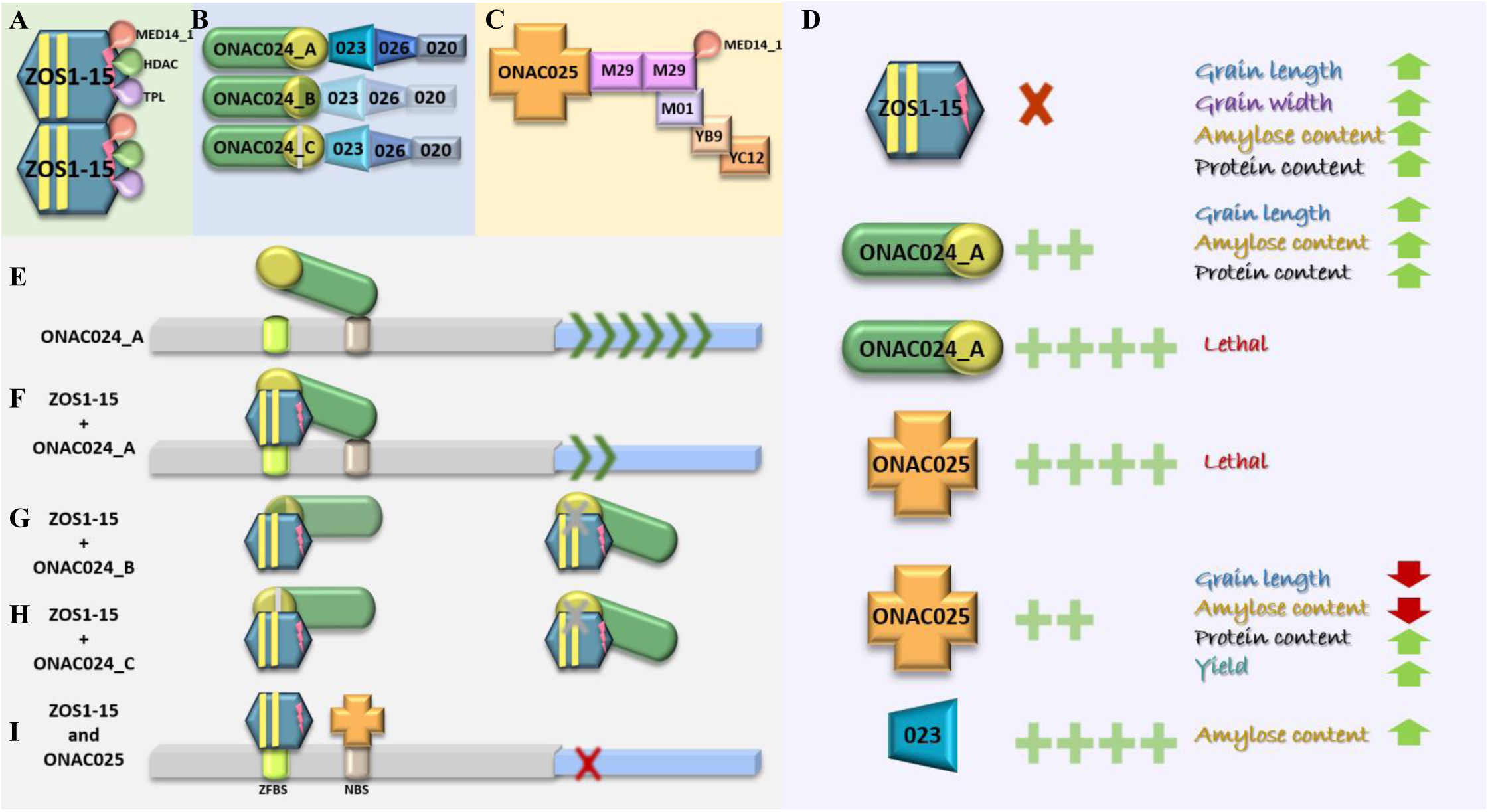
Model for a C2H2-NAC repressor-activator-competitor module controlling rice grain traits. **(A)** The zinc finger repressor complex is composed of ZOS1-15 (blue gray) dimers. ZOS1-15 has two C2H2 ZF motifs (yellow) and a DLN motif (pink). ZOS1-15 interacts with MED14_1, TOPLESS (TPL), histone deacetylases (HDACs). **(B)** The NAC activator complex comprises ONAC024 (green) and in tandem interactors ONAC023 (023; light blue), ONAC026 (026; dark blue) and ONAC020 (020; gray). ONAC024 is alternatively spliced. The isoforms (ONAC024_A, ONAC024_B and ONAC024_C) have a conserved NAC domain (green) and alterations in TRR (yellow). The color gradations indicate strength of interaction of isoforms of ONAC024 with ONAC023. **(C)** The NAC competitor complex involves ONAC025 (orange) which interacts with MADS29 (M29; pink). M29 homodimerizes and interacts with MED14_1. The in tandem interactors are MADS01 (M01), NF- YB9 (YB9) and NF-YC12 (YC12). The MADS-NF-Y complex has been constituted from our previous publications (Das et al., 2019; Malik et al., 2019) and available literature (Nayar et al., 2014). **(D)** The knock-down or knock-out of *ZOS1-15* leads to increased grain length, width, AAC and TPC. Regulated expression of *ONAC024* also leads to a similar grain phenotype. However, over expression of *ONAC024* is lethal. So is the over expression of *ONAC025*. However, the regulated expression of *ONAC025* decreases grain length and AAC, but increases TPC and grain yield. In addition, the over expression of *ONAC023* increases amylose content only. **(E)** Activator ONAC024_A binds to NAC binding site (NBS) in the promoters of downstream genes and activates them. **(F)** ZOS1-15 binds to ONAC024 and to the zinc finger binding site (ZFBS) in the promoter of downstream genes to equilibrate their expression. **(G, H)** The alternatively spliced isoforms ONAC024_B and ONAC024_C weakly bind to ZOS1-15, hence sequestering it and preventing its binding to ONAC024_A. **(I)** When the competitor ONAC025 replaces ONAC024 at NBS, the expression of downstream targets is affected.

In our study, the ZF repressor complex consists of a C_2_H_2_ ZF TF, ZOS1-15 with a DLN motif that homodimerizes and interacts with co-repressor TPL and HDT702 to execute its repressive action. It also interacts with a Mediator subunit MED14_1 (**Figure 10A**). C_2_H_2_ zinc finger TFs such as SUPERMAN, RABBIT EARS, ZFP207, ZCT1 harbor EAR motifs and function as transcriptional repressors (Li et al., 2016; Mortensen et al., 2019; Duan et al., 2021; Rodas et al., 2021). Such transcriptional repressors often interact with co-repressors TPL or TPRs and HDACs (Wang et al., 2020a; Zhuang et al., 2020; Hu et al., 2022b; Li et al., 2022a). Repressor ZOS1-15, interacts with a strong activator ONAC024. The activator complex consists of ONAC024, which interacts with ONAC023. ONAC023 interacts with ONAC026, which further interacts with ONAC020 (**Figure 10B**). Earlier, ONAC026 and ONAC020 have been reported to function as bifunctional TFs (Mathew et al., 2016; Wang et al., 2020b). NAC TFs often interact with other TFs to either enhance or secure their transcriptional activities (Harrington et al., 2019). TaNAC019 interacts with TaGAMyb to regulate glutelin and starch accumulation (Gao et al., 2021). Interaction of VvNAC26 and VvMADS9 is crucial for regulating seed and fruit development in grape (Zhang et al., 2021). OsCUC1 and OsCUC3 dimerize and function redundantly in regulating meristem/ organ boundary specification in rice (Wang et al., 2021). ONAC127 and ONAC129 form a heterodimer critical for rice grain filling wherein, ONAC129 negatively regulates the transcriptional activity of ONAC127 (Ren et al., 2021).

A complete knock-out of ZOS1-15 was lethal and its ectopic overexpression produced short statured plants as was the case when its interactor, ONAC024 was ectopically overexpressed, which caused highly stunted plant growth (**Figure 10D**). Any alteration in the expression of *ONAC024* had a negative effect on panicle architecture and grain filling. Grain filling was severely affected when ZOS1-15 expression was altered. The ectopic overexpression of MADS29 also resulted in severely stunted plants, thinner and smaller leaves, and failed grain filling (Nayar et al., 2013). The suppressed expression of MED14_1 caused reduced plant height, narrower leaves and negatively altered panicle morphology (Malik et al., 2020). The overexpression of ONAC023, an interactor of ONAC024, also had a negative effect on plant height, panicle architecture and days to flowering. Retarded growth of ONAC023_OE was also reported in another study (Li et al., 2022b). The ectopic overexpression of SS1/ ONAC025, the competitor for ONAC024 at NBS, was lethal, as reported earlier by us (Mathew et al., 2020).

The various grain traits controlled by the module components are grain length, width, AAC, TPC and grain yield (**Figure 10D**). Phenotyping data indicates that while ONAC024 has a positive influence over most traits, ZOS1-15 and ONAC025 behave in an opposite manner. However, ONAC025 enhances grain yield while ONAC023 only increases amylose content. An inverse relation has been often been reported between grain number and grain size. The knock-down of *RBG1* resulted in reduced grain size but increased grain yield (Lo et al., 2020). The overexpression of *WRKY70* resulted in increased grain size, but reduced grain yield (Tang et al., 2022). The overexpression plant of *GL6* showed increased grain length while, decreased tiller number, grains per panicle and yield (Wang et al., 2019). The knock-down and knock-out mutants of *OsGATA6* have early flowering, increased grain size but reduced grain number (Zhang et al., 2022). Since there is a considerable overlap between the traits controlled by these genes, the individuality of each component is a matter of speculation at this juncture. Amongst these traits, starch type is an integral part of grain quality, synthesized during endosperm development and influences rice grain size and its physiochemical properties. The starch biosynthetic pathway has been comprehensively studied in rice and the key enzymes have been elucidated. However, the transcriptional regulation of starch biosynthesis requires validation. This study indicates that ONAC024 promotes amylose production while, ZOS1-15 decreases it by oppositely regulating many starch biosynthesis genes. *GBSSI*, which promotes amylose production (Pérez et al., 2019) is up regulated in 024_SOE grains. So is *FLO7/ FLOURY ENDOPSERM 7* which regulates the development of amyloplast in the peripheral endosperm (Zhang et al., 2016). Similar pattern is seen for *BEIIb*. *BEIIb* deficiency results in higher proportion of amylopectin long chains, but shorter amylose chain length, resulting in higher gelatinization and chalkiness (Hu et al., 2022a). The 1-15_OE grain phenotype is similar to *ssiiia* mutants (Ryoo et al., 2007). This justifies the densely packed polyhedral shaped starch granules in the endosperm of 024_SOE. The relationship between chalkiness, starch structure and composition are well known. Increased chalky area or chalkiness rate significantly lowers amylose content (Deng et al., 2021). ONAC023 has also been characterized for its role in maintaining sugar homeostasis between source and sink organs (Li et al., 2022b). It has been reported that starch metabolism related genes are regulated by sugars (Hizukuri et al., 1989; Shen et al., 2022). Reduced expression of *MADS29* decreases AAC and results in spherical starch granules with large air spaces (Nayar et al., 2013). Size and number of endosperm cells and the physiochemical and biochemical properties of starch synthesis directly controls grain size, which affects weight (Fan et al., 2019). Generally, longer grains have higher amylose content (Bhardwaj et al., 2019), as was also seen here.

The other trait influenced is TPC, of which SSPs are the major components. Rice *SSPs* are specifically expressed during rice grain filling stages (Kawaksatsu et al., 2010) and maintain grain quality. SSP synthesis is spatially and temporarily controlled by specific and combinatorial binding of different TFs to the *cis*-elements of *SSP* promoters (Yang et al., 2023, Zhang et al., 2019). We identified ONAC024 as a positive regulator of TPC while, ZOS1-15 had a negative effect. This could be because of opposite regulation of various SSPs by them. ONAC024 bound to NBS and ZOS1-15 to ZFBS, which were present in the promoters of *GLU6* and *ALB13* (**Figure 10F**). As seen in our *in planta* reporter effector assays, the presence of both these TFs was able to decrease the activation of the *SSPs*. It can be hypothesized here that this interaction is able to balance the downstream gene expression, mitigating the effects of over excessive presence of ONAC024. It seems ONAC024 is an essential TF, and mechanisms have been put in place to equilibrate its effects, as its ectopic presence is harmful. Other members of this activator complex, OsNAC20 and OsNAC26 also regulate various *SSPs* (*GluA1*, *GluB4/5* and *16 KDa prolamin*) by binding to their promoters (Wang et al., 2020b).

In addition, ONAC024 exerts a feedback on ZOS1-15. Its alternatively spliced forms (**Figure 10B**), ONAC024_B and ONAC024_C do not bind to NBS. Their interaction with ZOS1-15 probably sequesters the repressor (**Figure 10G, H**). This will ensure more unhindered downstream activity by ONAC024. This is also in compliance with the reduced activity of ZOS1-15 (KD or KO), which results in a phenotype similar to 024_SOE. In terms of phenotype, both spliced forms have a positive effect on grain length and width. Both do not affect the AAC while only ONAC024_C affects TPC. Hence, a change in TRR domain, responsible for interaction with ONAC023, indicates a functional divergence amongst the spliced forms with respect to the main form. Such functional divergence amongst spliced variants has been shown for the two alternative splice variants of FLOWERING LOCUS M, FLM-β and FLM-δ, in regulating flowering at increased temperature. The FLM-β and FLM-δ variants contain the DNA-binding MADS-box domain. The FLM-δ does not bind to the DNA but competes with FLM-β isoform for its interaction with SHORT VEGETATIVE PHASE protein and binding to the promoters of *SEP3*, *SOC1* and *ATC* (Scortecci et al., 2001; Posé et al., 2013). The two alternatively spliced isoforms of Dlf1/ OsWRKY11, OsWRKY11.1 and OsWRKY11.2 regulate flowering and plant height differentially in rice (Cai et al., 2014). OsGS1:1a and OsGS1:1b, the two functional alternative spliced variants of OsGS1:1 differentially regulate nitrogen use efficiency (NUE), amylose content and grain development (Liu et al., 2022b).

Worth mentioning here is the delayed flowering phenotype of 1-15_OE plants. Increase in days to flowering could be explained by increased expression of *OsNF-YB9*, *OsNF-YC12*, *OsMADS5O*, *OsHd1* and lowered expression of *RFT1*. Overexpression of *NF-YB9* shows, increased plant height, delay in flowering, non-viable pollen and morphological defects in reproductive organs (Das et al., 2019). Overexpression of *Hd1* and reduced expression of *RFT1* causes delay in flowering (Pasriga et al., 2019). Members of the competitor complex discussed here, MADS29 (M29) and MADS01 (M01) are known to cause early flowering (Nayar et al., 2013). This shows that these genes can control other essential traits, apart from regulating grain phenotype.

Another repressor complex reported in this paper is composed of SS1/ ONAC025, which interacts with MADS29-NF-Y complex. The MADS-NF-Y complex, as reported in literature, includes MADS29 homodimer and the interactors of MADS29-MED14_1 and MADS01. M01 interacts with NF-YB9, which further interacts with NF-YC12 (YC12) (Das et al., 2019; Malik et al., 2020; Nayar et al., 2013) (**Figure 10C**). EMSA supported by contrasting grain phenotypes of 024_SOE and 025_SOE imply that ONAC025 competes with ONAC024 in binding to NBS on downstream promoters. Studies have reported competitions within TFs to regulate the same target genes. Repressor VvMYB30 and activator VvMYB14 compete for binding to the same *cis*-elements in the *VvSTS15/21* promoter to regulate stilbene biosynthesis (Mu et al., 2023). Activator EjMYB1 and repressor EjMYB2 regulate lignification by competitive binding with AC elements in the promoter of lignin biosynthesis genes in *Eriobotrya japonica* fruit (Xu et al., 2014). TFs ELK1 and GABP, compete to repress and activate the expression of *IPO4*, respectively (Xu et al., 2017). Competition by ONAC025 might be another mechanism to ensure regulated effects of ONAC024. The downstream genes common to be both ONAC024 and ONAC025 remain to be determined. These will be the ones whose equilibrated expression is essential for directly influencing various grain traits. In addition, ONAC025 has a positive effect on grain yield, which is not controlled by any other member of the complex.

Hence, the dynamic states of repressor-activator-competitor module discussed in this paper, comprising of ZOS1-15, ONAC024, ONAC023, ONAC026, ONAC020, SS1/ ONAC025 and MADS29, appear to be responsible for fine tuning grain length, width, AAC, TPC and grain yield.

## METHODS

### Plant material

*Oryza sativa indica* genotypes, IET-10364 and IR64, were grown in NIPGR field during Kharif season. Rice seed developmental stages have been divided into five categories named as, S1 (0-2 DAP), S2 (3-4 DAP), S3 (5-10 DAP), S4 (11-20 DAP) and S5 (21-29 DAP) (Agarwal et al., 2007; Agarwal et al., 2011). Rice seeds were tagged at the day of pollination (DAP) and were considered as zero DAP. Subsequently, the seeds for each DAP were tagged and harvested, dipped in liquid nitrogen and stored in -80 °C for further analysis. Panicles at two developmental growth stages P1 (0-5 cm) and P2 (5-10 cm) and other vegetative tissues like node, internode, leaf sheath, flag leaf, second leaf, root and callus (raised from *O*. *sativa indica* genotype, IET-10364 by culturing on callus induction N_6_B_5_ media for 12 days) were also harvested similarly.

### Sub-cellular localization studies

YFP fusion proteins were constructed by amplifying full length *ZOS1-15*, *ZOS1-15* truncated forms, full length *ONAC024* (ONAC024_A) and its isoforms (ONAC024_B and ONAC024_C) by gene specific primers (**Supplemental Table 1**) and cloned in pSITE-3CA (Chakrabarty et al., 2007) through Gateway^TM^ cloning technology (Invitrogen; USA). Vector pSITE 3CA has N-terminal YFP under the control of enhanced *cauliflower mosaic virus 35S* (*CaMV35S*) promoter. Truncated forms of ZOS1-15 were constructed by exclusion of N-terminal leucine rich region (LRR) forming ZOS1-15 without nucleolar retention signal (YFP-(ΔNoRS)-1-15), removal of C-terminal LRR forming ZOS1-15 without nuclear export signal (YFP-(ΔNES)-1-15) and removal of both N-terminal and C-terminal LRR forming YFP-(ΔNoRS/ΔNES)-1-15. The sub-cellular localization was observed in onion epidermal cells through particle bombardment using Biolistic-PDS-1000/He (Bio-Rad; USA) system (Casas et al., 1995). The YFP fused constructs were also transformed in *Agrobacterium* strain EHA105 (Xu and Qingshun, 2008) for transient expression in *N. benthamiana* leaf epidermal cells through *Agrobacterium* mediated infiltration (Norkunas et al., 2018). YFP fluorescence was observed under LEICA TCS SP-5 confocal microscope at 514 nm. The nuclear DNA specific fluorophore 4′,6-diamidino-2-phenylindole (DAPI) was visualized at 358 nm. Nuclear marker, *P_R82_* :*2* × *RFP* (Huang et al., 2014) and *AtWAK2*-pBIN2 endoplasmic reticulum (ER)-mCHERRY marker (He et al., 1999) both bearing RFP fluorescence were visualized at 588 nm.

### Confirmation of the alternatively spliced forms of O*NAC024* genes

The alternatively spliced forms of *ONAC024* were confirmed by semi-quantitative PCR amplification of the spliced fragments using specific and common primers flanking the region of variation amongst the isoforms (**Supplemental Table 1**). Amplification was carried out with Phusion^®^ high-fidelity Polymerase (New England Biolabs^®^*Inc*; USA) and the amplicons were separated by carrying out electrophoresis on 3 % ultrapure^TM^ (Invitrogen^TM^; USA) agarose gel. Amplified products were cloned in pJET1.2/blunt vector (ThermoScientific^TM^; USA) and were confirmed by sequencing. Alternate forms were translated by GeneRunner Version 3.05 to study the effect of these variations at the protein level.

The 1352 bp ORF region of *ONAC024* was amplified from genomic DNA of *indica* rice genotype IET-10364 and cloned through Gateway^TM^ cloning technology (Invitrogen; USA) in pSITE-3CA vector. The primers used for amplification are listed in **Supplemental Table 1**. The fusion protein was transiently expressed in *N. benthamiana* leaves through *Agrobacterium* mediated infiltration. Plants expressing YFP:ONAC024 and empty vector pSITE-3CA were harvested without mid vein after 48 hrs of *Agrobacterium* infiltration. Approximately 1 gm of harvested sample was grounded into a fine powder in mortar pestle with liquid nitrogen. To the grounded sample, 600 µL of extraction buffer (1M Tris buffer pH 8.0, 5 M NaCl, 0.5 M EDTA, 50% glycerol, 10% triton X100, 1M DTT, protease inhibitor cocktail, 50 mM PMSF) was added and the sample was homogenized. The solution was centrifuged at 13000 rpm for 15 mins at 4°C. The supernatant was filtered using mira cloth (Merck; USA). The total protein was electrophoresed on 10% PAGE gel (30% acrylamide mix, 1.5 M tris pH 8.8, 10% SDS, 10% APS, TEMED) followed with Western blot detection (Mahmood and Yang, 2012) using YFP antibody (Merck; USA).

### RNA isolation and qRT-PCR

Approximately 100 mg of *N. benthamiana* leaves were harvested for total RNA isolation using Trizol reagent (Ambion^®^; USA). RNA was isolated from S4 and S3 stage of transgenic and WT rice seeds for ZOS1-15 and ONAC024, respectively. Total RNA isolation with purification and quality check was performed as described previously (Agarwal et al., 2007). For making first stand cDNA from the purified RNA, high-capacity cDNA reverse transcriptase kit (ThermoScientific^TM^; USA) was used and the steps were followed as per its user manual. 2 µg of purified RNA was taken for making cDNA. The qRT-PCR was performed by using KAPA SYBR^®^ FAST qPCR Master Mix (2X) Universal with ROX Low Reference Dye (50X) (Sigma Aldrich^®^; USA). The primers used for qRT-PCR have been enlisted in **Supplemental Table 1**. The experiment was performed in two biological replicates and three technical replicates for each sample. The expression of each gene was normalized with rice *Actin* (*ACTI*) in rice seeds (Agarwal et al., 2007; Mathew et al., 2016). Fold change was calculated by 2-ΔΔ*Ct* method (Rao et al., 2013).

### Vector construction and generation of rice transgenic plants

For *ZOS1-15* overexpression construct (1-15_OE), full-length cDNA clone was obtained from KOME (Knowledge-based Oryza Molecular biological Encyclopedia, accession ID AK108227, http://cdna01.dna.affrc.go.jp/cDNA/). It was cloned into pB_4_NU vector (Raghuvanshi, 2001) between *Kpn*I and *BamH*I sites, downstream of maize *UBIQUITIN* promoter through pBSK+ vector. The full-length sequence of *ONAC024, ONAC023* and *SS1/ ONAC025* were taken from the rice genome annotation project (RGAP, http://rice.uga.edu/). For generating the overexpression construct of ONAC024 (024_OE), the full-length cDNA sequence of *ONAC024* was amplified from the S3 stage of IR64 grains using specific primers (**Supplemental Table 1**). The amplified cDNA was then cloned into the pB_4_NU vector between the *Sma*I and *BamH*I restriction sites. Overexpression construct of *ONAC023* (023_OE) were prepared by cloning ONAC023 between *Kpn*I and *BamH*I restriction sites in the binary vector, pB_4_NU. For construction of *ONAC024* seed-preferential overexpression construct, the CDS region was amplified with gene-specific primers (**Supplemental Table 1**) and cloned through Gateway^®^ cloning technology in pG_6_SOE vector containing the 2 kb promoter sequence of *GluD-1*. For generating the SS1/ ONAC025 seed-preferential overexpression construct, the full-length cDNA sequence of *SS1/ ONAC025* was amplified with specific primers (**Supplemental Table 1**) and cloned through Gateway^®^ cloning technology in pG_6_SOE vector.

To design the *ZOS1-15* RNAi (1-15_KD) and *ONAC024* RNAi (024_KD) constructs, a 159 bp full length coding sequence for *ZOS1-15* and 424 bp unique region for *ONAC024* were amplified along with CACC attached to forward primer (**Supplemental Table 1**) and cloned into Gateway entry vector pENTR^TM^/D-TOPO^®^ (ThermoScientific^TM^, USA) as per manufacturer’s protocol. The gene was cloned in pANDA vector in opposite orientation between two *attR* sites flanking the GUS linker gene (Reynolds et al., 2004) using Gateway^®^ based cloning. For *ONAC024* amiRNA construct preparation, the cDNA sequence was used as an input sequence in WMD3-Web MicroRNA Designer (http://wmd3.weigelworld.org/cgi-bin/webapp.cgi) with default parameters. The strategy used for amiRNA design (Warthmann et al., 2008) has been explained in **Supplemental Figure 18**. A 21mer miRNA partially complementary to the target mRNA was selected and appropriate miRNA* sequence was designed. miRNA* would pair to the miRNA and exactly mimic the stem-loop structure of the endogenous osa-MIR528 (Liu et al., 2005). miRNA construct was engineered using pNW55 vector (Addgene; USA), which contained the stem sequence of osa-MIR528 replaced by miRNA-miRNA* sequence by using four primers listed in **Supplemental Table 1** (024_I miR-s-FP has the miRNA in sense orientation, 024_II miR-a-RP is its reverse complement, 024_III miR*s-FP is the miRNA* sequence in sense and 024_IV miR*a-RP is the miRNA* sequence in the antisense orientation). Using an overlapping cloning procedure (Bryksin and Matsumura, 2010), three modification PCRs were performed by using ThermoScientific^TM^ Phusion^TM^ High-Fidelity DNA Polymerase kit. Each PCR reaction was carried out by using primer combinations G-4368-FP + 024_II miR-a-RP, 024_I miR-s-FP + 024_IV miR*a-RP and 024_III miR*s-FP + G-4369-RP on pNW55 vector as a template, and the resulting fragments of 256 bp, 87 bp and 259 bp length, respectively, were obtained. The three resulting fragments were gel purified (ThermoScientific^TM^ GeneJET Gel Extraction kit) according to the manufacturer’s protocol. For fusion PCR, 1µl mixture of purified PCR products were taken as a template in a 100 µl reaction with 20 µl 5X GC buffer, 10 µl dNTPs, 5 µl G-4368 -FP and 5 µl G-4369-RP (Warthmann et al., 2008). The final product of 554 bp was gel purified and cloned in pJET vector by using ThermoScientific^TM^ CloneJET PCR Cloning Kit. The positive plasmids were excised by *BamH*I and *Kpn*I restriction enzymes and further cloned in pB_4_NU binary vector.

To generate mutants of *ZOS1-15* by CRISPR analysis, we designed a single gRNA targeting the first ZF domain and another gRNA targeting the DLN motif within *ZOS1-15* CDS sequence using CRISPR-GE tool (http://skl.scau.edu.cn/targetdesign/) (Xie et al., 2017). A unique stretch of 20 nt gRNA fused with tRNA was synthesized by Sigma-Aldrich^®^ to produce gRNA-tRNA sequence. Primers used to form gRNA-tRNA construct are listed in **Supplemental Table 1**. pGTR vector was used to amplify gRNA-tRNA sequence. The cloning of gRNA-tRNA sequence in pRGEB32 vector (Xie et al., 2015) was done using Phusion^TM^ High-Fidelity PCR kit (ThermoScientific^TM^, USA). The first reaction involved the amplification of gRNA-tRNA sequence (127 bp) using gRNA forward primer and L3AD5 reverse primer. The second reaction involved the amplification of gRNA-tRNA sequence using L5AD5 forward primer and gRNA reverse primer. Bands from both reactions were eluted using GeneJet Gel extraction kit (ThermoScientific^TM^, USA) and an overlapping PCR was carried out. Overlapping PCR included two reactions, the former reaction involved annealing of 20 ng of elute each from first and second reaction using Phusion polymerase (ThermoScientific^TM^, USA). The first reaction was paused after 10 cycles and second reaction mixture (with L5AD5 forward and L3AD5 reverse primers) was added. Following PCR, the obtained 244 bp fragment band was digested with *Fok*I enzyme. *Fok*I digested gRNA-tRNA sequence was cloned in the pRGEB32 vector (**Supplemental Figure 2C**; Addgene, USA) using Golden gate kit (ThermoScientific^TM^). The positive clones were confirmed by sequencing.

To distinguish amongst the roles of ONAC024 and its isoforms, a Chimeric REpressor gene Silencing Technology (CRES-T) was employed (Mitsuda et al., 2011). The CDS region of ONAC024 and its isoforms were amplified with gene specific forward primer and the reverse primers contained an EAR motif (Singh et al., 2019). A chimeric repressor produced by a fusion of 024_A, 024_B and 024_C with the repressor motif generated 024_A_CRES-T, 024_B_CRES-T and 024_C_CRES-T constructs. The positive clones were confirmed by restriction digestion and sequencing.

All confirmed clones for all the eleven constructs generated in this study, were transformed in *Agrobacterium* strain EHA105 and through *Agrobacterium* mediated transformation were incorporated into the rice genotype IET-10364 using a scutellum-transformation based protocol (Toki et al., 2006). All the rice transgenic plants were selected on 50 μg/mL hygromycin.

### Screening of transgenic plants

1-15_OE, 024_SOE, 024_A_CRES-T, 024_B_CRES-T, 024_C_CRES-T, 023_OE and 025_SOE transgenic plants were screened for *GUS* activity through histochemical *GUS* staining (Jefferson et al., 1987). A small piece of leaf from transgenic plants was incubated in GUS buffer, 0.5 mM potassium ferricyanide, 0.5 mM potassium ferrocyanide, 0.1% (v/v) Triton-X-100, 50 mM phosphate buffer (pH 7.0), 10 mM sodium EDTA and 1 mg/ml (w/v) X-gluc (5-bromo-4-chloro-3-indolyl-β-D-glucuronide) containing 10% methanol at 37 °C for 16-20 hours. Blue colour of the leaves indicated GUS activity, which was visualized under Nikon AZ100 microscope. All the *ZOS1-15, ONAC024, ONAC023* and *SS1/ ONAC025* transgenic plants were screened for the presence of *HYGROMYCIN PHOSPHOTRANSFERASEII*/*HPTII* gene (850 bp) using the genomic DNA of transgenic plants. The primers used for screening transgenic plants are listed in **Supplemental Table 1**.

1-15_KO (generated by CRISPR-Cas9 technology) was screened for the presence of indels in the target region of *ZOS1-15* in the T_0_ generation and in the subsequent T_1_ generations. Genomic DNA was extracted from all the plants (Dellaporta et al., 1983). The target region of *ZOS1-15* was amplified using flanking primers specific to the coding sequence of *ZOS1-15*. The amplified product was cloned in pJET vector using CloneJET PCR Cloning Kit (ThermoScientific^TM^, USA). Eight positive clones from each line were given for DNA sequencing. The raw sequence data were aligned to coding sequence of *ZOS1-15* using Clustal Omega tools (https://www.ebi.ac.uk/Tools/msa/clustalo/). In the subsequent T_1_ generation, plants were screened for presence of *HPTII* gene using hygromycin specific primers. To find T-DNA free plants, *Cas9* amplification (549 bp) was performed with Cas9 specific primers and target sequence amplification (321 bp) was performed with vector specific primers (*U3* promoter) using genomic DNA as a template. The primers used for screening transgenic plants are listed in **Supplemental Table 1**.

### Detection of off-target mutations induced by CRISPR Cas9 effector

CRISPR-GE tool was used to identify any off-target editing derived from the target sequence. The off-target genes were chosen on the basis of maximum off-target score for the detection of off-target mutations. Genomic DNA was extracted from selected lines of transgenic plants. Target region of off-target genes were amplified using gene-specific flanking primers and genomic DNA as template (**Supplemental Table 1**). The amplified product from plants of selected lines, was cloned in pJET vector using CloneJET PCR Cloning Kit (ThermoScientific^TM^, USA). Eight positive clones from each plant were given for Sanger sequencing. The raw sequence data were aligned to coding sequence of *ZOS1-15* using Clustal Omega tools (https://www.ebi.ac.uk/Tools/msa/clustalo/).

### Phenotypic analysis of transgenic rice plants

Phenotypic analysis of transgenic and wild type (WT) rice plants was done by measuring agronomic traits like seed length, seed width and seed weight using WinSEEDLE^TM^ software (Regent Instrument Inc.; Canada). Images of plants and seeds were taken using a Nikon-5200 DSLR camera. For scanning electron microscopy (SEM, EVO^®^ LS10 Zeiss, Germany), mature grains of *ZOS1-15 ONAC024* and *SS1/ ONAC025* transgenic plants and wild type grains were examined. SEM was used to examine cell size in outer epidermal layer of lemma cells. Mid-central part of mature grains was examined under 500X and 1X magnification. Images were analyzed using ImageJ (Schneider et al., 2012). For the measurement of lemma cell size and number in longitudinal and transverse direction, only those values were considered in there is significant change in both the lines. To study the morphology of starch granules, mature grains of *ZOS1-15*, *ONAC024, ONAC023* and *SS1/ ONAC025* transgenic plants along with wild-type grains were dehusked and sectioned longitudinally with the help of a scalpel. Central parts of endosperm were examined at 2500X magnification. The compound starch granules were marked (Zhang et al., 2016). Mature grains of *ZOS1-15*, *ONAC024, ONAC023* and *SS1/ ONAC025* transgenic plants were harvested and used to study grain chalkiness. Hand-cut transverse sections of dehusked grains were observed under 2.5X magnification in a stereozoom microscope (AZ100, Nikon; Japan).

### Tissue sectioning and microscopic studies

Leaf samples from WT and OE_024 plants were fixed with FAE (10 % formaldehyde, 5 % acetic acid and 50 % absolute ethanol) fixative, and dehydrated with ethanol. The samples were paraplast X-TRA® (Sigma-Aldrich; USA) embedded, sliced on a rotary microtome (Leica Biosystems, Germany). The sections stained with 0.25 % toluidine blue-O/TBO were analyzed and captured under stereozoom microscope (AZ100, Nikon; Japan) and light microscope (Eclipse 80i, Nikon; Japan).

### Amylose estimation in rice seeds

Dehusked mature rice seeds (100 mg) were ground into a fine powder with the help of liquid nitrogen in a mortar-pestle. Total amylose content in rice seed was measured by iodine binding affinity (Juliano B, 1971). The fine powder was transferred into a 100 ml volumetric flask, and 1 ml of 95% ethanol and 9 ml of 1 M NaOH was added. The solution was boiled for 30 mins at 100 °C to gelatinize starch. Then the flask was kept at room temperature for 10 mins and adequate distilled water was added to make up the volume to 100 ml. 5 ml of this starch solution was added to another 100 ml volumetric flask and 1 ml of 1 M acetic acid and 2 ml of iodine solution (0.2 gm of iodine and 2 gm of potassium iodide in 100 ml MQ) were added into it and mixed well. The solution was left to rest for 30 mins. Following this, a colorimetric assay was performed by taking the absorbance at 620 nm in the spectrophotometer (ThermoScientific^TM^, USA). The amylose content was calculated based on a standard graph (Avaro et al., 2011).

### Total protein isolation from rice seeds

Mature rice seeds from S5 stage (200 mg) were ground to a fine powder in a mortar and pestle using liquid nitrogen. The powder was homogenized in 600 µL of protein extraction buffer (1 M Tris pH 7.5, 5 M NaCl, 0.5 M EDTA, 50 mM DTT, 100 mM PMSF). The homogenate was incubated on ice for 15 mins and then centrifuged at 13000 rpm for 10 mins at 4 °C (Lang et al., 2013). The supernatant containing total seed protein was transferred to a fresh MCT and separated on a 10% SDS-PAGE. Total protein concentrations of each of these samples were identified using Bradford assay (Bradford, 1976). Absorbance of the protein sample was measured at 595 nm.

### Auto-activation test by yeast two-hybrid assays

To check the transcriptional auto-activation of ONAC024, multiple deletion constructs named as, A, AB, C, D, DE, ABCD, ABCDE, E+TRR, TRR_A, TRR_B, TRR_C were constructed carrying ONAC024 domains or sub-domains and named accordingly. The truncated fragments were amplified from ONAC024 pENTR clone and cloned in pGBKT7-BD vector (Takara^TM^ Bio Inc; USA) through restriction-based cloning. For checking the transcriptional auto-activation activity of ONAC024 construct fused with DLN motif (024_A_CRES-T, 024_B_CRES-T and 024_C_CRES-T), the fragments were cloned in pGBKT7-BD vector (Addgene; USA). The primer sequences are listed in **Supplemental Table 1**. The reconstituted plasmids had truncated fragments fused with GAL4 DNA binding domain of pGBKT7. The constructs were transformed in yeast strain AH109. Reconstituted rGAL4 and pGBKT7-BD vectors were taken as positive and negative controls for the experiment, respectively. The yeast transformation was performed by using Quick and Easy Yeast Transformation Mix Kit (Takara^TM^ Bio Inc; USA) and steps were as per its user manual. The transformed yeast cells were selected by plating onto synthetic drop-out (SD) medium lacking tryptophan and SD/-Trp/-His/-Ade (TDO) media with 10 mM 3-amino-1,2,4-triazole (3-AT) in comparison with relevant controls. To quantify the auto-activation of ONAC024 and its isoforms fused with DLN motif, β-Galactosidase Assay was performed as per Clonetech^®^ yeast protocol handbook.

### Protein interactions in yeast by yeast two-hybrid assays

Yeast two-hybrid assays were carried out using the Matchmaker Gold system (Takara™ Bio USA, Inc) with yeast strain AH109. Truncated forms of ZOS1-15 were developed by removing the N-terminal ZF domain, C-terminal ZF domain, and C-terminal DLN motif, thus, forming ZOS1-15_ΔN, ZOS1-15_ΔC and ZOS1-15_ΔDLN constructs, respectively. These constructs were cloned in pENTR™ D-TOPO^®^ vectors and recombined into Gateway™ compatible vectors, pGADT7-GW and pGBKT7-GW (Addgene; USA), respectively, using Gateway™ LR Clonase™ II Enzyme mix (Invitrogen™; USA). Full length coding sequence of ZOS1-15, MED14_1, and HDACs (HDT701, HD704, HD706, HD709, HD711, SRT701) were cloned in pGADT7 and pGBKT7 vectors (Takara™ Bio USA, Inc) using restriction-based cloning (primers listed in **Supplemental Table 1**). Further, HDT702, OsTPL, ONAC024, ONAC023, ONAC020, ONAC026, SS1/ ONAC025 and MADS29 CDS were cloned in pGADT7-GW and pGBKT7-GW vectors using Gateway™-based cloning. The alternative spliced isoforms of ONAC024 (ONAC024_B and ONAC024_C) and its deletion constructs (NAC, TRR_A, TRR_B and TRR_C) were cloned in pGBKT7-GW vectors using Gateway™-based cloning. All the primers are listed in **Supplemental Table 1**. The pGADT7 - pGBKT7 or pGADT7-GW and pGBKT7-GW plasmid constructs were co-transformed into AH109 yeast strain using EZ-Yeast^TM^ transformation kit (MP Biomedicals; USA) and plated on synthetic drop-out (SD) medium lacking leucine and tryptophan. Positive interactors were detected by incubating transformed yeast cells on SD/-Leu/-Trp/-His/-Ade and SD/-Leu/-Trp/-His/-Ade with 80 mg/L X-α-gal (5-bromo-4-chloro-3-indolyl-β-D-galactopyranoside) medium as mentioned previously (Mathew et al., 2016). To inhibit the auto-activation ability of ONAC020 and ONAC026, the yeast media containing SD/-AHLT was supplemented with 10 mM 3-Amino-1,2,4-triazole (3-AT). Co-transformed yeast cells with T-antigen_pGADT7 and p53_pGBKT7 were used as a positive control while co-transformed yeast cells with T-antigen_pGADT7 and Lam_pGBKT7 were used as a negative control.

### Protein expression and *in vitro* pull-down assays

Full-length coding sequences of ZOS1-15, MED14_1, ONAC023, ONAC026, ONAC020 and MADS29 were cloned in GST (Glutathione-S-transferase) tagged vector, pGEX4T-1 (Addgene; USA) and ZOS1*-*15, HDT702, OsTPL, ONAC024, ONAC023 ONAC026 and SS1/ ONAC025 CDS were cloned in MBP (maltose binding protein) tagged vector, pMALc2x (Addgene; USA) by restriction-based cloning to produce GST_ZOS1-15, GST_MED14_1, GST_ONAC023, GST_ONAC026, GST_ONAC020, GST_MADS29, MBP_ZOS1-15, MBP_HDT702, MBP_OsTPL, MBP_LC2, MBP_ONAC024, MBP_ONAC023, MBP_ONAC026 and MBP_SS1/ ONAC025 protein expression constructs, respectively. Primers used in this study are listed in **Supplemental Table 1**. Proteins were expressed in competent *Escherichia coli* strain BL21 by adding 1 mM IPTG (isopropyl-β-D-thiogalactopyranoside) at 37°C for 4-5 hours. MBP-fused proteins were purified using MBP resin and the GST-tagged proteins were purified with glutathione agarose beads (G-biosciences^®^; USA). For *in vitro* pull-down assays, GST-fused protein or GST (control) was bound to glutathione agarose beads (G-biosciences^®^; USA), overnight, with gentle rotation, followed by 2-3 times washing with 1X PBS (0.14 M NaCl, 2.7 mM KCl, 10.1 mM Na_2_HPO_4_, and 1.8 mM KH_2_PO_4_). Subsequently, MBP fused proteins were added and incubated for 4 hrs. Final washing was done with wash buffer containing 30 mM Tris–HCl, 50 mM NaCl, pH 7.5. Protein complex was eluted using 2.8 mg/ml reduced glutathione (Sigma-Aldrich^®^; USA). Alternatively, MBP-fused proteins or MBP (control) were bound to MBP resin (G-biosciences^®^; USA). These were incubated with GST-fused protein and similar steps were performed as mentioned above. The protein complex was eluted using 3 mg/ml maltose (Sigma-Aldrich^®^; USA). Eluted proteins were denatured and separated on SDS-PAGE gel and analyzed by immunoblotting with anti-GST antibody (1:4000; G7781-Sigma-Aldrich^®^) and anti-MBP antibody (1:4000; M6295-Sigma-Aldrich^®^). Empty pGEX4T-1 and empty pMAL-c2x were used as a control in all interactions.

### BiFC assay

Full length cDNAs of *ZOS1-15*, *HDT702*, O*NAC024*, *ONAC023, SS1/ ONAC025* and *MADS29* were PCR amplified using gene-specific primers (listed in **Supplemental Table 1**) and cloned into the Gateway^TM^ entry vector, pENTR^TM^/D-TOPO^®^ (Invitrogen^TM^; USA). pENTR constructs were then transferred by an LR reaction into the destination vector, cd3-1648/pSITE-nEYFP-C1 and cd3-1651/pSITE-cEYFP-N1 (Martin et al., 2009) in-frame to the N-terminal half or C-terminal half of YFP, respectively, to form YFP^N^-ZOS1-15, ZOS1-15-YFP^C^, YFP^N^-HDT702, HDT702-YFP^C^, YFP^N^-ONAC024, ONAC024-YFP^C^, YFP^N^-ONAC023, ONAC023-YFP^C^, YFP^N^-SS1/ ONAC025, SS1/ ONAC025-YFP^C^ and YFP^N^-MADS29, MADS29-YFP^C^ constructs. These constructs were co-transfected into leaf cells of *N. benthamiana* by *Agrobacterium*-mediated infiltration (Zhang et al., 2020). Empty vector constructs were used as control. Following two days of incubation at 28°C after infection, leaf tissue was observed under LEICA TCS-SP5 Confocal Laser Scanning Microscope for YFP signal at 514 nm.

### Co-immunoprecipitation assay

For Co-IP assay, ZOS1-15, ONAC024, SS1/ ONAC025 were cloned in pSITE-3CA vector in-frame with N-terminal YFP tag and ZOS1-15, MED14_1, HDT702, TPL, ONAC023, MADS29 were cloned in pGWB420 vector in-frame with C-terminal MYC tag. These constructs were transformed in *Agrobacterium* strain EHA105 for infiltration in three-week-old *N. benthamiana* leaves. In control, empty YFP tag vector with MYC plasmids were co-transfected whereas in case of test, YFP_ZOS1-15 or YFP_ONAC024 or YFP_SS1/ ONAC025 with MYC plasmids (MED14_1/HDT702/TPL/ZOS1-15/ ONAC023/ MADS29_MYC) were co-transfected in equal molar ratio. The infiltrated plants were kept at 28°C in dark for 48 hrs and the 100 mg of leaf samples were then harvested. Total protein from leaf samples were harvested using 3-4 ml of protein extraction buffer (20 mM Tris-Cl, pH, 7.5, 150 mM NaCl, 1mM EDTA, 1mM DTT and 1mM PMSF). The cell lysate were centrifuged at 100g for 10 min at 4°C. Co-IP assay was performed using Dynabeads™ Protein A Immunoprecipitation Kit (ThermoScientific^TM^, USA). The supernatant were separated and incubated with 50 μL of Dynabeads Protein A (pre-equilibrated with 1-3 µg of anti-MYC antibody) for 4 h at 4°C and washed three times with wash buffer and finally eluted using 20 µl elution buffer (given in kit). These samples were then analyzed by immunoblotting with anti-YFP antibody (1:2500; G7781-Sigma-Aldrich^®^) and anti-MYC antibody (1:10000; M6295-Abcam).

### Yeast one-hybrid assay

For yeast one-hybrid assay, the upstream 2 kb promoter sequence of *ProGLU6* and *ProALB13* were amplified from the genomic DNA of *indica* rice genotype IR64. The promoters were cloned in the vector pABAi (Addgene; USA) and the primer sequences are listed in **Supplemental Table 1**. To generate the deletion constructs of *ProGLU6*, five fragments of 421 bp each, including a 21 bp overlap were amplified from *ProGLU6*_pABAi construct and named as *ProGLU6_1*, *ProGLU6_2*, *ProGLU6_3*, *ProGLU6_4* and *ProGLU6_5*. Similarly, the deletion fragments of *ProALB13* were constructed and named accordingly. These acted as the bait constructs. For the prey construct, CDS region of ZOS1-15, ONAC024 and SS1/ ONAC025 were amplified using gene-specific primers (**Supplemental Table 1**) and cloned in pGADT7-GW/pGADT7-AD (Addgene, USA) through Gateway^®^ cloning technology. These were named as ZOS1-15_AD, ONAC024_AD and SS1/ ONAC025_AD. Promoters cloned in pABAi were transformed in yeast strain YIH Gold using the Quick and Easy Yeast Transformation Mix Kit (Takara^TM^ Bio Inc; USA). The transformed yeast cells were selected by plating onto synthetic drop-out (SD) medium (0.667% yeast nitrogen base, 2% glucose and appropriate auxotrophic amino acid supplements) lacking uracil. The minimal inhibitory concentration of the bait vector was estimated by spotting the yeast colonies transformed with the promoter constructs in pABAi vector on SD/-Ura supplemented with 100-800 ng/mL Aureobasidin. The growth of the bait vector was stopped at 200 ng/mL Aureobasidin. Further, in an overnight grown culture of the bait strain growing in SD/-Ura broth, ZOS1-15_AD or ONAC024_AD or SS1/ ONAC025 _AD was transformed. The transformed yeast cells were selected by plating onto SD medium lacking uracil, leucine and supplemented with 200 ng/mL Aureobasidin. pABAi vector co-transformed with ZOS1-15_AD or ONAC024_AD or SS1/ ONAC025 _AD was taken as the negative control.

### Electrophoretic mobility shift assay (EMSA)

The promoter sequences were checked for known ZF binding site (ZF_BS) and NAC binding site (NBS) in PLANT PAN3.0 (http://plantpan.itps.ncku.edu.tw/). ZF_BS was present in *ProGLU6_1* and *ProALB13_3* deletion fragments and NBS was present in *ProGLU6_4* and *ProALB13_3* deletion fragments of the 2 kb promoter sequence of *GLU6* and *ALB13*, respectively. The ZF_BS for *ProGLU6* and *ProALB13* was CACT. The NBS for *ProGLU6* was TTACT and CACGTAA for *ProALB13*. For synthesizing the EMSA probes (Sigma Aldrich^®^; US), the above-mentioned binding sites along with its flanking base-pairs from the promoter sequence were taken in multiples of two or three, 2X ZF_BS for *ProGLU6* and *ProALB13*, 2X NBS for *ProGLU6* and 3XNBS for *ProALB13*. The binding site sequences were also mutated maintaining the flanking sequences. The mutated binding site was GTTC for mZF_BS. The mNBS sequences were GCGAG for *ProGLU6* and AGTCAGG for *ProALB13*. The probes were labelled using Biotin 3’ end DNA labelling kit (Invitrogen^TM^; US). The efficiency of the labelled probes was estimated by dot plot assay as mentioned in the user manual. GST_ZOS1-15, MBP_ONAC024 and MBP_SS1/ ONAC025 fusion proteins were used for EMSA. The protein inductions were carried out with 0.5 mM IPTG and the bacterial incubation conditions were 28°C with 200 rpm for 8 hrs followed by cell lysis through sonication and protein purification. EMSA was performed using LightShift^®^ chemiluminscent EMSA kit (ThermoScientific^TM^; USA). The binding reactions were prepared along with appropriate controls. Individually, 30 µg of GST_ZOS1-15 or MBP_ONAC024 or MBP_SS1/ ONAC025 purified proteins were incubated with 25 fmol of labelled 2X_ZF_BS/ m2X_ZF_BS and 2X_NBS/ m2X_NBS, respectively. Two different concentrations of unlabeled probes (50X, 100X) were used to check the specificity of binding. The binding reactions were kept at RT for 20 mins. For competitive EMSA assay the concentration of MBP_SS1/ ONAC025 purified protein was increased from 50X to 100X while, keeping the concentration of MBP_ONAC024 and probe constant. However, later the concentration of MBP_NAC04 was raised from 50X to 100X without changing the concentration of MBP_SS1/ ONAC025 and probe. A native polyacrylamide (PAGE) gel (6%) was prepared in 0.5X TBE for resolving the protein probe complex. The biotin-labelled DNA probes were detected using the Chemiluminescent Nucleic Acid Detection Module Kit (ThermoScientific^TM^; USA) and pictured under BioRad Gel Doc^TM^ XR (BioRad; USA).

### Reporter effector assay

For reporter effector assay, CDS region of ZOS1-15, ONAC024, MADS29 MADS29+ZOS1-15_DLN and SS1/ ONAC025 were cloned in pEARLYGate201 (Addgene; USA) (**Supplemental Table 1**). These effector plasmids had ZOS1-15/ ONAC024/ MADS29/ MADS29+ZOS1-15_DLN/ SS1/ ONAC025 overexpressed under the control of *CaMV35S* promoter. The 2 kb promoter sequences of *GLU6*, *ALB13* and *Cys-Prt* were cloned in pMDC164 vector (Curtis and Grossniklaus, 2003) using Gateway^®^ cloning technology. These reporter plasmids contained *ProGLU6*, *ProALB13* and *ProCys-Prt* in-frame with reporter gene *β-glucuronidase* (*GUS*). The plasmids were transformed in *Agrobacterium* strain EHA105 for in-filtration in *N. benthamiana* leaves. In control, empty effector plasmid with reporter vector was transfected whereas in test, and effector and reporter constructs both were injected in equal molar ratio. To check the repression activity of ZOS1-15_DLN/ ZOS1-15 on MADS29/ ONAC024 and SS1/ ONAC025 on ONAC024 both constructs in equal molar ratio, along with reporter construct were used. The infiltrated plants were kept at 28°C in dark for 48 hrs and then the leaf samples were harvested. The harvested samples were used for RNA isolation and expression analysis by qRT-PCR. The reporter expression was normalized with *HPTII*. The control was the expression of *GUS* in the presence of empty effector vector, pEARLYGate201. The expression of *Basta* (*blpR*) and MADS29/ *ZOS1-15/ ONAC024/ SS1/ ONAC025* was examined to confirm adequate effector activity. The experiment was performed in six biological replicates for each sample, with three technical replicates. The GraphPad Prism v7 (https://www.graphpad.com/scientific-software/prism/) was used to make dot-whisker plot.

### Statistical analysis

All the experiments were performed in biological replicates and statistical significance was calculated using one-tailed student t-test assuming equal variance for two samples in Microsoft Excel^®^. The *p<0.05, **p<0.01 and ***p<0.005 were considered as significant.

## ACCESSION NUMBERS

Sequence data for this article can be found in the Rice Genome Annotation Project (http://rice.uga.edu/) under the following LOCUS IDs: *ALB13* (LOC_Os07g11650), *GLU6* (LOC_Os02g15090), *HD704* (LOC_Os07g06980), *HD706* (LOC_Os06g37420), *HD709* (LOC_Os11g09370), *HD711* (LOC_Os04g33480), *HDT701* (LOC_Os05g51830), *HDT702* (LOC_Os01g68104), *MADS29* (LOC_Os02g07430), *MED14_1* (LOC_Os08g24400), *ONAC020* (LOC_Os01g01470), *ONAC023* (LOC_Os02g12310), *ONAC024* (LOC_Os05g34310), *ONAC025* (LOC_Os05g31330), *ONAC026* (LOC_Os01g29840), *SRT701* (LOC_Os04g20270), *TPL* (LOC_Os08g06480), and *ZOS1-15* (LOC_Os01g62190).

## Supplemental Data

**Supplemental Figure 1.** Expression of *ZOS1-15* in transgenic plants.

**Supplemental Figure 2.** Generation of *ZOS1-15* knock-out mutant rice plants using CRISPR-Cas9 technology.

**Supplemental Figure 3.** Off-target analysis in the CRISPR-Cas9 generated knock-out lines of *ZOS1-15*.

**Supplemental Figure 4**. Cell size and cell number in the husk of *ZOS1-15* transgenic and mutant grains.

**Supplemental Figure 5.** Phenotypic analysis of *ZOS1-15* transgenic and mutant plants.

**Supplemental Figure 6.** Panicle morphology and yield related traits of *ZOS1-15* transgenic and mutant plants.

**Supplemental Figure 7.** Regulation of flowering by ZOS1-15.

**Supplemental Figure 8.** Protein dimerizing with ZOS1-15.

**Supplemental Figure 9.** Yeast two-hybrid assays of ONAC024 spliced variants and their deletion constructs with ZOS1-15.

**Supplemental Figure 10.** Expression and subcellular localization of ZOS1-15 and ONAC024

**Supplemental Figure 11.** Sub-cellular localization of YFP:ZOS1-15 and YFP:ONAC024_A/B/C.

**Supplemental Figure 12.** Protein structure and SDR-mediated alternative splicing of *ONAC024*.

**Supplemental Figure 13.** The transcriptional activity of ONAC024.

**Supplemental Figure 14.** Expression analysis in rice transgenic plants overexpressing *ONAC024.*

**Supplemental Figure 15.** Phenotypic analysis of rice transgenic plants overexpressing *ONAC024.*

**Supplemental Figure 16.** Leaf morphology of 024_OE plants.

**Supplemental Figure 17.** PCR protocol to synthesize amiRNA construct for ONAC024 from pNW55 vector.

**Supplemental Figure 18.** The transcriptional activity of ONAC024 isoforms fused with DLN motif.

**Supplemental Figure 19.** Expression of *ONAC024* in transgenic plants.

**Supplemental Figure 20.** Cell size and number in husk of *ONAC024* transgenic grains.

**Supplemental Figure 21.** Vegetative morphology of 024_SOE rice plants.

**Supplemental Figure 22.** Panicle phenotyping of *ONAC024* transgenic plants.

**Supplemental Figure 23.** Grain chalkiness of *ZOS1-15* and *ONAC024* transgenic and mutant grains.

**Supplemental Figure 24.** Differential expression of starch biosynthesis related genes in *ZOS1-15* and *ONAC024* transgenic and mutant grains.

**Supplemental Figure 25.** Expression of *SSPs* and its regulation by ZOS1-15 and ONAC024.

**Supplemental Figure 26.** Interaction of ONAC024 and ONAC023.

**Supplemental Figure 27.** Phenotyping of *ONAC023* over expression lines.

**Supplemental Figure 28.** Total protein content in 023_OE lines compared to WT.

**Supplemental Figure 29.** Interaction of ONAC023 with ONAC026 and of ONAC026 with ONAC020.

**Supplemental Figure 30.** Yeast two-hybrid assay to check the interactions of transcription factors belonging to ZOS-NAC module.

**Supplemental Figure 31.** SS1/ ONAC025 binds to the promoters of *GLU6* and *ALB13*.

**Supplemental Figure 32.** Reporter effector assay to confirm binding of SS1/ ONAC025 to *ProGLU6* and *ProALB13*.

**Supplemental Figure 33.** SS1/ ONAC025 interferes with ONAC024 for regulating the expression of *SSP* encoding genes.

**Supplemental Figure 34.** Phenotypic analysis of 025_SOE rice plants.

**Supplemental Figure 35.** Cell size and number in husk of 025_SOE transgenic grains.

**Supplemental Figure 36.** Regulation of grain starch by *SS1/ ONAC025*.

**Supplemental Figure 37.** *SS1/ ONAC025* regulates the accumulation of SSPs.

**Supplemental Figure 38.** Yeast two-hybrid assay to check the interactions of SS1/ ONAC025 with other members of ZOS-NAC module.

**Supplemental Figure 39.** Interaction between SS1/ ONAC025 and MADS29.

**Supplemental Table 1.** Primers used in this study.

## AUTHORS CONTRIBUTIONS

P.A. designed, supervised the experiments and arranged for funding. P.J., A.M., F.Q. and A.V. performed experiments related to ZOS1-15. R.P., U.D., I.E.M., A.P. and A.V. performed experiments related to ONAC024. I.E.M., P.J., A.P.V., A.Y. and U.D. performed experiments related to ONAC023. A.K.T. guided with important scientific inputs during investigation and manuscript writing. P.A., P.J., R.P., A.Y., A.P.V., A.M., UD and F.Q. analysed the data, prepared figures, and wrote the article. All authors have discussed and approved the article.

## Acknowledgments

This work was supported by research grant numbers BT/AB/NIPGR/SEED BIOLOGY/2012 by Department of Biotechnology (DBT) and CRG/2018/000501 and SPF/2021/002899 by Science and Engineering Research Board (SERB) under the Ministry of Science and Technology, India. The authors acknowledge the core grant from National Institute of Plant Genome Research (NIPGR), New Delhi for funding this work and central instrumentation (CIF) facilities of NIPGR for scanning electron microscopy, stereomicroscopy and confocal microscopy experiments. We thank the NBT e-library consortium for providing online access to research articles. R.P., I.E.M., A.M., A.Y., A.P.V., U.D and A.P., acknowledge University Grants Commission for JRF and SRF fellowships. P.J., F.Q and A.V. are thankful to Council of Scientific and Industrial Research and Department of Biotechnology, respectively, for JRF and SRF fellowships.

## References

Agarwal, P., Kapoor, S., and Tyagi, A.K. (2011). Transcription factors regulating the progression of monocot and dicot seed development. BioEssays 33, 189–202.

Agarwal, P., Arora, R., Ray, S., Singh, A.K., Singh, V.P., Takatsuji, H., Kapoor, S., and Tyagi, A.K. (2007). Genome-wide identification of C2H2 zinc-finger gene family in rice and their phylogeny and expression analysis. Plant Mol. Biol. 65, 467–485.

Avaro, M.R.A., Pan, Z., Yoshida, T., and Wada, Y. (2011). Two alternative methods to predict amylose content of rice grain by using tristimulus CIE lab values and developing a specific color board of starch-iodine complex solution. Plant Prod. Sci. 14, 164–168.

Bhardwaj, M., Sandhu, K.S., and Saxena, D.C. (2019). Experimental and modeling studies of the flow, dynamic and creep recovery properties of pearl millet starch as affected by concentration and cultivar type. Int. J. Biol. Macromol. 135, 544–552.

Bradford, M.M. (1976). A rapid and sensitive method for the quantitation of microgram quantities of protein utilizing the principle of protein-dye binding. Anal. Biochem. 72, 248–254.

Bryksin, A.V., and Matsumura, I. (2010). Overlap extension PCR cloning: a simple and reliable way to create recombinant plasmids. BioTechniques 48, 463–465.

Cai, Y., Chen, X., Xie, K., Xing, Q., Wu, Y., Li, J., Du, C., Sun, Z., and Guo, Z. (2014). Dlf1, a WRKY transcription factor, is involved in the control of flowering time and plant height in rice. PLoS ONE 9(7): e102529

Casas, A.M., Kononowicz, A.K., Bressan, R.A., and Hasegawa, P.M. (1995). Cereal transformation through particle bombardment. Plant Breed Rev. 13, 235–264.

Chakrabarty, R., Banerjee, R., Chung, S.M., Farman, M., Citovsky, V., Hogenhout, S.A., Tzfira, T., and Goodin, M. (2007). PSITE vectors for stable integration or transient expression of autofluorescent protein fusions in plants: probing *Nicotiana benthamiana*-virus interactions. Mol. Plant Microbe Interact. 20, 740–750.

Chen, W., Chen, L., Zhang, X., Yang, N., Guo, J., Wang, M., Ji, S., Zhao, X., Yin, P., Cai, L., Xu, J., Zhang, L., Han, Y., Xiao, Y., Xu, G., Wang, Y., Wang, S., Wu, S., Yang, F., Jackson, D., Cheng, J., Chen, S., Sun, C., Qin, F., Tian, F., Fernie, A.R., Li, J., Yan, J., and Yang, X. (2022). Convergent selection of a WD40 protein that enhances grain yield in maize and rice. Science (N.Y.) 375, 6585.

Cheng, X., Zhang, S., Tao, W., Zhang, X., Liu, J., Sun, J., Zhang, H., Pu, L., Huang, R., and Chen, T. (2018). INDETERMINATE SPIKELET1 recruits histone deacetylase and a transcriptional repression complex to regulate rice salt tolerance. Plant Physiol. 178, 824–837.

Christianson, J.A., Dennis, E.S., Llewellyn, D.J., and Wilson, I.W. (2010). ATAF NAC transcription factors: regulators of plant stress signaling. Plant Signal Behav. 5, 428–432.

Cordenunsi-Lysenko, B.R., Nascimento, J.R.O., Castro-Alves, V.C., Purgatto, E., Fabi, J.P., and Peroni-Okyta, F.H.G. (2019). The starch is (not) just another brick in the wall: the primary metabolism of sugars during banana ripening. Front. Plant Sci. 10, 391.

Curtis, M.D., and Grossniklaus, U. (2003). A gateway cloning vector set for high-throughput functional analysis of genes *in planta*. Plant Physiol. 133, 462–469.

Das, S., Parida, S.K., Agarwal, P., and Tyagi, A.K. (2019). Transcription factor OsNF-YB9 regulates reproductive growth and development in rice. Planta 250, 1849–1865.

Dellaporta, S.L., Wood, J., and Hicks, J.B. (1983). A plant DNA minipreparation: Version II. Plant Mol. Biol. Rep. 1, 19–21.

Deng, F., Li, Q., Chen, H., Zeng, Y., Li, B., Zhong, X., Wang, L., and Ren, W. (2021). Relationship between chalkiness and the structural and thermal properties of rice starch after shading during grain-filling stage. Carbohydr. Polym. 252, 117212.

Deng, H., Chen, Y., Liu, Z., Liu, Z., Shu, P., Wang, R., Hao, Y., Su, D., Pirrello, J., Liu, Y., Li, Z., Grierson, D., Giovannoni, J.J., Bouzayen, M., and Liu, M. (2022). SlERF.F12 modulates the transition to ripening in tomato fruit by recruiting the co-repressor TOPLESS and histone deacetylases to repress key ripening genes. Plant Cell 34, 1250–1272.

Diaz, I., Vicente-Carbajosa J Fau-Abraham, Z., Abraham Z Fau-Martínez, M., Martínez M Fau-Isabel-La Moneda, I., Isabel-La Moneda I Fau-Carbonero, P., and Carbonero, P. (2002) The GAMYB protein from barley interacts with the DOF transcription factor BPBF and activates endosperm-specific genes during seed development. Plant J. 29(4):453–64

Dong, S., Dong, X., Han, X., Zhang, F., Zhu, Y., Xin, X., Wang, Y., Hu, Y., Yuan, D., Wang, J., Huang, Z., Niu, F., Hu, Z., Yan, P., Cao, L., He, H., Fu, J., Xin, Y., Tan, Y., Mao, B., Zhao, B., Yang, J., Yuan, L., and Luo, X. (2021). OsPDCD5 negatively regulates plant architecture and grain yield in rice. Proc. Natl. Acad. Sci. U S A 118, 29.

Dong, X., Zhang, D., Liu, J., Liu, Q.Q., Liu, H., Tian, L., Jiang, L., and Qu le, Q. (2015). *Plastidial disproportionating* enzyme participates in starch synthesis in rice endosperm by transferring maltooligosyl groups from amylose and amylopectin to amylopectin. Plant Physiol. 169, 2496–2512.

Duan, M., Ke, X.J., Lan, H.X., Yuan, X., Huang, P., Xu, E.S., Gao, X.Y., Wang, R.Q., Tang, H.J., Zhang, H.S., and Huang, J. (2021). A Cys2/His2 zinc finger protein acts as a repressor of the green revolution gene *SD1/OsGA20ox2* in rice (*Oryza sativa L*.). Plant Cell Physiol. 61, 2055–2066.

Fan, C., Wang, G., Wang, Y., Zhang, R., Wang, Y., Feng, S., Luo, K., and Peng, L. (2019). Sucrose synthase enhances hull size and grain weight by regulating cell division and starch accumulation in transgenic rice. Int. J. Mol. Sci. 20, 4971

Feng, B.H., Han, Y.C., Xiao, Y.Y., Kuang, J.F., Fan, Z.Q., Chen, J.Y., and Lu, W.J. (2016). The banana fruit Dof transcription factor MaDof23 acts as a repressor and interacts with MaERF9 in regulating ripening-related genes. J. Exp. Bot. 67, 2263–2275.

Gao, Y., An, K., Guo, W., Chen, Y., Zhang, R., Zhang, X., Chang, S., Rossi, V., Jin, F., Cao, X., Xin, M., Peng, H., Hu, Z., Guo, W., Du, J., Ni, Z., Sun, Q., and Yao, Y. (2021). The endosperm-specific transcription factor TaNAC019 regulates glutenin and starch accumulation and its elite allele improves wheat grain quality. Plant Cell 33, 603–622.

Han, Y.-Y., Zhou, H.-Y., Xu, L.-A., Liu, X.-Y., Fan, S.-X., and Cao, J.-S. (2018). The zinc-finger transcription factor BcMF20 and its orthologs in Cruciferae which are required for pollen development. Biochem. Biophys. Res. Commun. 503, 998–1003.

Hao, D., Ohme-Takagi M Fau-Sarai, A., and Sarai, A. (1998) Unique mode of GCC box recognition by the DNA-binding domain of ethylene-responsive element-binding factor (ERF domain) in plant. J. Biol. Chem. 273(41):26857–61

Harrington, S.A., Overend, L.E., Cobo, N., Borrill, P., and Uauy, C. (2019). Conserved residues in the wheat (*Triticum aestivum*) NAM-A1 NAC domain are required for protein binding and when mutated lead to delayed peduncle and flag leaf senescence. BMC Plant Biol. 19, 407.

He, Z.H., Cheeseman, I., He, D., and Kohorn, B.D. (1999). A cluster of five cell wall-associated receptor kinase genes, Wak1-5, are expressed in specific organs of *Arabidopsis*. Plant Mol. Biol. 39, 1189-1196.

Hizukuri, S., Takeda, Y., Maruta, N., and Juliano, B.O. (1989). Molecular structures of rice starch. Carbohydr. Res. 189, 227–235.

Hoogenkamp, H., Kumagai, H., and Wanasundara, J. (2017). Rice protein and rice protein products. Sustainable protein sources (Elsevier), pp. 47–65.

Hori, K., and Sun, J. (2022). Rice grain size and quality. Rice (N.Y.) 15, 33.

Hu, Y., Zhang, Y., Yu, S., Deng, G., Dai, G., and Bao, J. (2022a). Combined effects of *BEIIb* and *SSIIa* alleles on amylose contents, starch fine structures and physicochemical properties of *indica* rice. Foods 12, 119.

Hu, Y., Han, Z., Sun, Y., Wang, S., Wang, T., Wang, Y., Xu, K., Zhang, X., Xu, X., Han, Z., and Wu, T. (2020). ERF4 affects fruit firmness through TPL4 by reducing ethylene production. Plant J. 103, 937–950.

Hu, Y., Han, Z., Wang, T., Li, H., Li, Q., Wang, S., Tian, J., Wang, Y., Zhang, X., Xu, X., Han, Z., Lü, P., and Wu, T. (2022b). Ethylene response factor MdERF4 and histone deacetylase MdHDA19 suppress apple fruit ripening through histone deacetylation of ripening-related genes. Plant Physiol. 188, 2166–2181.

Huang, D., Wang, S., Zhang, B., Shang-Guan, K., Shi, Y., Zhang, D., Liu, X., Wu, K., Xu, Z., Fu, X., and Zhou, Y. (2015). A gibberellin-mediated DELLA-NAC signaling cascade regulates cellulose synthesis in rice. Plant Cell 27, 1681–1696.

Huang, L., Hua, K., Xu, R., Zeng, D., Wang, R., Dong, G., Zhang, G., Lu, X., Fang, N., Wang, D., Duan, P., Zhang, B., Liu, Z., Li, N., Luo, Y., Qian, Q., Yao, S., and Li, Y. (2021). The LARGE2-APO1/APO2 regulatory module controls panicle size and grain number in rice. Plant Cell 33, 1212–1228.

Huang, Y.Y., Shi, Y., Lei, Y., Li, Y., Fan, J., Xu, Y.J., Ma, X.F., Zhao, J.Q., Xiao, S., and Wang, W.M. (2014). Functional identification of multiple nucleocytoplasmic trafficking signals in the broad-spectrum resistance protein RPW8.2. Planta 239, 455–468.

Jefferson, R.A., Kavanagh, T.A., and Bevan, M.W. (1987). GUS fusions: beta-glucuronidase as a sensitive and versatile gene fusion marker in higher plants. EMBO. J. 6, 3901–3907.

Jia, D., Jiang, Z., Fu, H., Chen, L., Liao, G., He, Y., Huang, C., and Xu, X. (2021). Genome-wide identification and comprehensive analysis of NAC family genes involved in fruit development in kiwifruit (*Actinidia*). BMC Plant Biol. 21, 44.

Jin, S.-K., Zhang, M.-Q., Leng, Y.-J., Xu, L.-N., Jia, S.-W., Wang, S.-L., Song, T., Wang, R.-A., Yang, Q.-Q., Tao, T., Cai, X.-L., and Gao, J.-P. (2022). *OsNAC129* regulates seed development and plant growth and participates in the brassinosteroid signaling pathway. Front. Plant Sci. 13, 905148.

Juliano B, O. (1971). A simplified assay for milled rice amylose. Cereal Sci. Today 16, 334–360.

Kawakatsu, T., Hirose, S., Yasuda, H., and Takaiwa, F. (2010). Reducing rice seed storage protein accumulation leads to changes in nutrient quality and storage organelle formation. Plant Physiol. 154, 1842–1854.

Kou, X., Zhou, J., Wu, C.E., Yang, S., Liu, Y., Chai, L., and Xue, Z. (2021). The interplay between ABA/ethylene and NAC TFs in tomato fruit ripening: a review. Plant Mol. Biol. 106, 223–238.

Kunieda, T., Mitsuda, N., Ohme-Takagi, M., Takeda, S., Aida, M., Tasaka, M., Kondo, M., Nishimura, M., and Hara-Nishimura, I. (2008). NAC family proteins NARS1/NAC2 and NARS2/NAM in the outer integument regulate embryogenesis in *Arabidopsis*. Plant Cell 20, 2631–2642.

Lang, G.H., Kagiya, Y., Ohnishi-Kameyama, M., and Kitta, K. (2013). Evaluation of extraction solutions for biochemical analyses of the proteins in rice grains. Biosci. Biotechnol. Biochem. 77, 126–131.

Laubscher, M., Brown, K., Tonfack, L.B., Myburg, A.A., Mizrachi, E., and Hussey, S.G. (2018). Temporal analysis of *Arabidopsis* genes activated by *Eucalyptus grandis* NAC transcription factors associated with xylem fibre and vessel development. Sci. Rep. 8, 10983.

Li, J., Wang, Y., Zhang, Y., Wang, W., Irish, V.F., and Huang, T. (2016). RABBIT EARS regulates the transcription of TCP4 during petal development in *Arabidopsis*. J. Exp. Bot. 67, 6473–6480.

Li, Y., Lei, R., Pu, M., Cai, Y., Lu, C., Li, Z., and Liang, G. (2022a). bHLH11 inhibits bHLH IVc proteins by recruiting the TOPLESS/TOPLESS-RELATED corepressors. Plant Physiol. 188, 1335–1349.

Li, Z., Wei, X., Tong, X., Zhao, J., Liu, X., Wang, H., Tang, L., Shu, Y., Li, G., and Wang, Y. (2022b). The OsNAC23-Tre6P-SnRK1a feed-forward loop regulates sugar homeostasis and grain yield in rice. Mol. Plant 15, 706–722.

Liu, B., Li, P., Li, X., Liu, C., Cao, S., Chu, C., and Cao, X. (2005). Loss of function of *OsDCL1* affects microRNA accumulation and causes developmental defects in rice. Plant Physiol. 139, 296–305.

Liu, G.S., Li, H.L., Grierson, D., and Fu, D.Q. (2022a). NAC transcription factor family regulation of fruit ripening and quality: A review. Cells 11, 525.

Liu, X., Tian, Y., Chi, W., Zhang, H., Yu, J., Chen, G., Wu, W., Jiang, X., Wang, S., and Lin, Z. (2022b). Alternative splicing of *OsGS1; 1* affects nitrogen-use efficiency, grain development, and amylose content in rice. Plant J. 110, 1751–1762.

Liu, Y., Khan, A.R., Azhar, W., Wong, C.E., Li, Y., Huang, Y., Cao, X., Liu, Z., and Gan, Y. (2022c). Cys2/His2-type zinc finger proteins regulate plant growth and development. Crit. Rev. Plant Sci. 41, 351–363.

Lo, S.F., Cheng, M.L., Hsing, Y.C., Chen, Y.S., Lee, K.W., Hong, Y.F., Hsiao, Y., Hsiao, A.S., Chen, P.J., Wong, L.I., Chen, N.C., Reuzeau, C., Ho, T.D., and Yu, S.M. (2020). Rice *Big Grain 1* promotes cell division to enhance organ development, stress tolerance and grain yield. Plant Biotechnol. J. 18, 1969–1983.

Lu, C., Mi, L.Z., Grey, M.J., Zhu, J., Graef, E., Yokoyama, S., and Springer, T.A. (2010). Structural evidence for loose linkage between ligand binding and kinase activation in the epidermal growth factor receptor. Mol. Cell Biol. 30, 5432–5443.

Lyu, T., and Cao, J. (2018). Cys₂/His₂ zinc-finger proteins in transcriptional regulation of flower development. Int. J. Mol. Sci. 19, 2589.

Ma, L., Li, R., Ma, L., Song, N., Xu, Z., and Wu, J. (2021). Involvement of NAC transcription factor NaNAC29 in *Alternaria alternata* resistance and leaf senescence in *Nicotiana attenuata*. Plant Divers. 43, 502–509.

Mahmood, T., and Yang, P.C. (2012). Western blot: technique, theory, and trouble shooting. N. Am. J. Med. Sci. 4, 429–434.

Malik, N., Ranjan, R., Parida, S.K., Agarwal, P., and Tyagi, A.K. (2020). Mediator subunit *OsMED14* plays an important role in rice development. Plant J. 101, 1411–1429.

Martin, K., Kopperud, K., Chakrabarty, R., Banerjee, R., Brooks, R., and Goodin, M.M. (2009). Transient expression in *Nicotiana benthamiana* fluorescent marker lines provides enhanced definition of protein localization, movement and interactions *in planta*. Plant J. 59, 150–162.

Mathew, I.E., and Agarwal, P. (2018). May the fittest protein evolve: favoring the plant-specific origin and expansion of NAC transcription factors. BioEssays 40, e1800018.

Mathew, I.E., Das, S., Mahto, A., and Agarwal, P. (2016). Three rice NAC transcription factors heteromerize and are associated with seed size. Front. Plant Sci. 7, 1638.

Mathew, I.E., Priyadarshini, R., Mahto, A., Jaiswal, P., Parida, S.K., and Agarwal, P. (2020). SUPER STARCHY1/ONAC025 participates in rice grain filling. Plant direct 4, e00249.

McCarty, A.S., Kleiger, G., Eisenberg, D., and Smale, S.T. (2003). Selective dimerization of a C2H2 zinc finger subfamily. Mol. Cell 11, 459–470.

Mendes, G.C., Reis, P.A., Calil, I.P., Carvalho, H.H., Aragão, F.J., and Fontes, E.P. (2013). GmNAC30 and GmNAC81 integrate the endoplasmic reticulum stress-and osmotic stress-induced cell death responses through a vacuolar processing enzyme. Proc. Natl. Acad. Sci. U S A 110, 19627–19632.

Mitsuda, N., Matsui, K., Ikeda, M., Nakata, M., Oshima, Y., Nagatoshi, Y., and Ohme-Takagi, M. (2011). CRES-T, an effective gene silencing system utilizing chimeric repressors. Methods Mol. Biol. 754, 87–105.

Mortensen, S., Weaver, J.D., Sathitloetsakun, S., Cole, L.F., Rizvi, N.F., Cram, E.J., and Lee-Parsons, C.W.T. (2019). The regulation of ZCT1, a transcriptional repressor of monoterpenoid indole alkaloid biosynthetic genes in Catharanthus roseus. Plant Direct 3, e00193.

Mu, H., Li, Y., Yuan, L., Jiang, J., Wei, Y., Duan, W., Fan, P., Li, S., Liang, Z., and Wang, L. (2023). MYB30 and MYB14 form a repressor-activator module with WRKY8 that controls stilbene biosynthesis in grapevine. Plant Cell 35, 552–573.

Nan, J., Feng, X., Wang, C., Zhang, X., Wang, R., Liu, J., Yuan, Q., Jiang, G., and Lin, S. (2018). Improving rice grain length through updating the GS3 locus of an elite variety Kongyu 131. Rice 11, 21.

Nayar, S., Sharma, R., Tyagi, AK., and Kapoor, S. (2013) Functional delineation of rice *MADS29* reveals its role in embryo and endosperm development by affecting hormone homeostasis. J. Exp. Bot. 64, 4239–4253

Nishioka, S., Sakamoto, T., and Matsunaga, S. (2020). Roles of BRAHMA and its interacting partners in plant chromatin remodeling. Cytologia 85, 263–267.

Niu, X., Luo, D., Gao, S., Ren, G., Chang, L., Zhou, Y., Luo, X., Li, Y., Hou, P., Tang, W., Lu, B.-R., and Liu, Y. (2010). A conserved unusual posttranscriptional processing mediated by short, direct repeated (SDR) sequences in plants. J. Genet. Genomics 37, 85–99.

Norkunas, K., Harding, R., Dale, J., and Dugdale, B. (2018). Improving agroinfiltration-based transient gene expression in *Nicotiana benthamiana*. Plant Methods 14, 71.

Nuruzzaman, M., Manimekalai, R., Sharoni, A.M., Satoh, K., Kondoh, H., Ooka, H., and Kikuchi, S. (2010). Genome-wide analysis of NAC transcription factor family in rice. Gene 465, 30–44.

Ohtani, H., and Iwasaki, Y.W. (2021). Rewiring of chromatin state and gene expression by transposable elements. Dev. Growth Differ. 63, 262–273.

Pasriga, R., Yoon, J., Cho, L.-H., and An, G. (2019). Overexpression of *RICE FLOWERING LOCUS T 1 (RFT1)* induces extremely early flowering in rice. Mol. Cells 42, 406–417.

Pérez, L., Soto, E., Farré, G., Juanos, J., Villorbina, G., Bassie, L., Medina, V., Serrato, A.J., Sahrawy, M., Rojas, J.A., Romagosa, I., Muñoz, P., Zhu, C., and Christou, P. (2019). CRISPR/Cas9 mutations in the rice *Waxy/GBSSI* gene induce allele-specific and zygosity-dependent feedback effects on endosperm starch biosynthesis. Plant Cell. Rep. 38, 417–433.

Plant, A.R., Larrieu, A., and Causier, B. (2021). Repressor for hire! The vital roles of TOPLESS-mediated transcriptional repression in plants. New Phytol. 231, 963–973.

Posé, D., Verhage, L., Ott, F., Yant, L., Mathieu, J., Angenent, G.C., Immink, R.G., and Schmid, M. (2013). Temperature-dependent regulation of flowering by antagonistic FLM variants. Nature 503, 414–417.

Puentes-Romero, A.C., González, S.A., González-Villanueva, E., Figueroa, C.R., and Ruiz-Lara, S. (2022). AtZAT4, a C2H2-type zinc finger transcription factor from Arabidopsis thaliana, is involved in pollen and seed development. Plants 11, 1974.

Raghuvanshi, S. (2001). Investigations on chloroplast transformation and characterization of constitutively photomorphogenic 1 (COP1) gene in rice. Ph.D. Thesis, University of Delhi.

Rao, X., Huang, X., Zhou, Z., and Lin, X. (2013). An improvement of the 2^(-delta delta CT) method for quantitative real-time polymerase chain reaction data analysis. Biostat. Bioinforma. Biomath. 3, 71–85.

Ren, Y., Huang, Z., Jiang, H., Wang, Z., Wu, F., Xiong, Y., and Yao, J. (2021). A heat stress responsive NAC transcription factor heterodimer plays key roles in rice grain filling. J. Exp. Bot. 72, 2947–2964.

Reynolds, A., Leake, D., Boese, Q., Scaringe, S., Marshall, W.S., and Khvorova, A. (2004). Rational siRNA design for RNA interference. Nat. Biotechnol 22, 326–330.

Reynolds, N., O’Shaughnessy, A., and Hendrich, B. (2013). Transcriptional repressors: multifaceted regulators of gene expression. Dev. 140, 505–512.

Rodas, A.L., Roque, E., Hamza, R., Gómez-Mena, C., Minguet, E.G., Wen, J., Mysore, K.S., Beltrán, J.P., and Cañas, L.A. (2021). MtSUPERMAN plays a key role in compound inflorescence and flower development in *Medicago truncatula*. Plant J. 105, 816–830.

Ryoo, N., Yu, C., Park, C.S., Baik, M.Y., Park, I.M., Cho, M.H., Bhoo, S.H., An, G., Hahn, T.R., and Jeon, J.S. (2007). Knockout of a starch synthase gene *OsSSIIIa/Flo5* causes white-core floury endosperm in rice (*Oryza sativa* L.). Plant Cell Rep. 26, 1083–1095.

Schneider, C.A., Rasband, W.S., and Eliceiri, K.W. (2012). NIH Image to ImageJ: 25 years of image analysis. Nat. Methods 9, 671–675.

Schumann, U., Prestele, J., O’Geen, H., Brueggeman, R., Wanner, G., and Gietl, C. (2007). Requirement of the C3HC4 zinc RING finger of the *Arabidopsis* PEX10 for photorespiration and leaf peroxisome contact with chloroplasts. Proc. Natl. Acad. Sci. U S A 104, 1069–1074.

Scortecci, K.C., Michaels, S.D., and Amasino, R.M. (2001). Identification of a MADS-box gene, FLOWERING LOCUS M, that represses flowering. Plant J. 26, 229–236.

Sharma, R., Agarwal, P., Ray, S., Deveshwar, P., Sharma, P., Sharma, N., Nijhawan, A., Jain, M., Singh, A.K., Singh, V.P., Khurana, J.P., Tyagi, A.K., and Kapoor, S. (2012). Expression dynamics of metabolic and regulatory components across stages of panicle and seed development in indica rice. Funct. Integr. Genomics 12, 229–248.

Shen, L., Li, J., and Li, Y. (2022). Resistant starch formation in rice: Genetic regulation and beyond. Plant Commun. 3: 100329

Shende, R., Pandey, S., and Bhan, KS. (2019) Seed protein profiling of rice genotypes. J. Pharmacogn. Phytochem. 8, 4131–4137

Shi, C.-L., Dong, N.-Q., Guo, T., Ye, W.-W., Shan, J.-X., and Lin, H.-X. (2020). A quantitative trait locus GW6 controls rice grain size and yield through the gibberellin pathway. Plant J. 103, 1174–1188.

Singh, P., Mathew, I.E., Verma, A., Tyagi, A.K., and Agarwal, P. (2019). Analysis of rice proteins with DLN repressor motif/s. Int. J. Mol. Sci. 20, 1600.

Singh, S., Koyama, H., Bhati, K.K., and Alok, A. (2021). The biotechnological importance of the plant-specific NAC transcription factor family in crop improvement. J. Plant Res. 134, 475–495.

Takatsuji, H., Mori, M., Benfey, P.N., Ren, L., and Chua, N.H. (1992). Characterization of a zinc finger DNA-binding protein expressed specifically in *Petunia* petals and seedlings. EMBO. J. 11, 241–249.

Tang, J., Mei, E., He, M., Bu, Q., and Tian, X. (2022). Functions of OsWRKY24, OsWRKY70 and OsWRKY53 in regulating grain size in rice. Planta 255, 92.

Toki, S., Hara, N., Ono, K., Onodera, H., Tagiri, A., Oka, S., and Tanaka, H. (2006). Early infection of scutellum tissue with *Agrobacterium* allows high-speed transformation of rice. Plant J. 47, 969–976.

Tran, L.S., Nakashima, K., Sakuma, Y., Osakabe, Y., Qin, F., Simpson, S.D., Maruyama, K., Fujita, Y., Shinozaki, K., and Yamaguchi-Shinozaki, K. (2007). Co-expression of the stress-inducible zinc finger homeodomain ZFHD1 and NAC transcription factors enhances expression of the ERD1 gene in *Arabidopsis*. Plant J. 49, 46–63.

Trupkin, S.A., Astigueta, F.H., Baigorria, A.H., García, M.N., Delfosse, V.C., González, S.A., Pérez de la Torre, M.C., Moschen, S., Lía, V.V., Fernández, P., and Heinz, R.A. (2019). Identification and expression analysis of NAC transcription factors potentially involved in leaf and petal senescence in *Petunia hybrida*. Plant Sci. 287, 110195.

Vargas-Hernández, B.Y., Núñez-Muñoz, L., Calderón-Pérez, B., Xoconostle-Cázares, B., and Ruiz-Medrano, R. (2022). The NAC transcription factor ANAC087 induces aerial rosette development and leaf senescence in *Arabidopsis*. Front. Plant Sci. 13, 818107.

Verma, A., Prakash, G., Ranjan, R., Tyagi, A.K., and Agarwal, P. (2020). Silencing of an ubiquitin ligase increases grain width and weight in indica rice. Front. Genet. 11, 600378.

Wang, A., Hou, Q., Si, L., Huang, X., Luo, J., Lu, D., Zhu, J., Shangguan, Y., Miao, J., and Xie, Y. (2019). The PLATZ transcription factor GL6 affects grain length and number in rice. Plant Phys. 180, 2077–2090.

Wang, H., Jiao, X., Kong, X., Liu, Y., Chen, X., Fang, R., and Yan, Y. (2020a). The histone deacetylase HDA703 interacts with OsBZR1 to regulate rice brassinosteroid signaling, growth and heading date through repression of *Ghd7* expression. Plant J. 104, 447–459.

Wang, J., Chen, Z., Zhang, Q., Meng, S., and Wei, C. (2020b). The NAC transcription factors OsNAC20 and OsNAC26 regulate starch and storage protein synthesis. Plant Physiol. 184, 1775–1791.

Wang, J., Bao, J., Zhou, B., Li, M., Li, X., and Jin, J. (2021). The osa-miR164 target *OsCUC1* functions redundantly with *OsCUC3* in controlling rice meristem/organ boundary specification. New Phytol. 229, 1566–1581.

Wang, S., Wu, K., Yuan, Q., Liu, X., Liu, Z., Lin, X., Zeng, R., Zhu, H., Dong, G., Qian, Q., Zhang, G., and Fu, X. (2012). Control of grain size, shape and quality by *OsSPL16* in rice. Nat. Genet. 44, 950–954.

Wang, S., Li, S., Liu, Q., Wu, K., Zhang, J., Wang, S., Wang, Y., Chen, X., Zhang, Y., Gao, C., Wang, F., Huang, H., and Fu, X. (2015). The OsSPL16-GW7 regulatory module determines grain shape and simultaneously improves rice yield and grain quality. Nat. Genet. 47, 949–954.

Warthmann, N., Chen, H., Ossowski, S., Weigel, D., and Hervé, P. (2008). Highly specific gene silencing by artificial miRNAs in rice. PloS ONE 3, e1829.

Waters, B.M., Uauy, C., Dubcovsky, J., and Grusak, M.A. (2009). Wheat (*Triticum aestivum*) NAM proteins regulate the translocation of iron, zinc, and nitrogen compounds from vegetative tissues to grain. J. Exp. Bot. 60, 4263–4274.

Weng, J., Gu, S., Wan, X., Gao, H., Guo, T., Su, N., Lei, C., Zhang, X., Cheng, Z., Guo, X., Wang, J., Jiang, L., Zhai, H., and Wan, J. (2008). Isolation and initial characterization of GW5, a major QTL associated with rice grain width and weight. Cell Res. 18, 1199–1209.

Wu, Q., Liu, Y., Xie, Z., Yu, B., Sun, Y., and Huang, J. (2022). *OsNAC016* regulates plant architecture and drought tolerance by interacting with the kinases GSK2 and SAPK8. Plant Physiol. 189, 1296–1313.

Xie, K., Minkenberg, B., and Yang, Y. (2015). Boosting CRISPR/Cas9 multiplex editing capability with the endogenous tRNA-processing system. Proc. Natl. Acad. Sci. U S A 112, 3570–3575.

Xie, X., Ma, X., Zhu, Q., Zeng, D., Li, G., and Liu, Y.G. (2017). CRISPR-GE: A convenient software toolkit for CRISPR-based genome editing. Mol. Plant 10, 1246–1249.

Xu, P., Lin, W., Liu, F., Tartakoff, A., and Tao, T. (2017). Competitive regulation of IPO4 transcription by *ELK1* and *GABP*. Gene 613, 30–38.

Xu, Q., Yin, X.-r., Zeng, J.-k., Ge, H., Song, M., Xu, C.-j., Li, X., Ferguson, I.B., and Chen, K.-s. (2014). Activator-and repressor-type MYB transcription factors are involved in chilling injury induced flesh lignification in loquat via their interactions with the phenylpropanoid pathway. J. Exp. Bot. 65, 4349–4359.

Xu, R., and Qingshun, L.Q. (2008). Protocol: Streamline cloning of genes into binary vectors in Agrobacterium via the Gateway® TOPO vector system. Plant Methods 4, 4.

Xu, Y., Prunet, N., Gan, E.S., Wang, Y., Stewart, D., Wellmer, F., Huang, J., Yamaguchi, N., Tatsumi, Y., Kojima, M., Kiba, T., Sakakibara, H., Jack, T.P., Meyerowitz, E.M., and Ito, T. (2018). SUPERMAN regulates floral whorl boundaries through control of auxin biosynthesis. EMBO. J. 37.

Xun, Q., Mei, M., Song, Y., Rong, C., Liu, J., Zhong, T., Ding, Y., and Ding, C. (2022). SWI2/SNF2 chromatin remodeling ATPases SPLAYED and BRAHMA control embryo development in rice. Plant Cell Rep. 41, 1389–1401.

Yamamoto, T., Yoshida, Y., Nakajima, K., Tominaga, M., Gyohda, A., Suzuki, H., Okamoto, T., Nishimura, T., Yokotani, N., and Minami, E. (2018). Expression of RSOsPR10 in rice roots is antagonistically regulated by jasmonate/ethylene and salicylic acid via the activator OsERF87 and the repressor OsWRKY76, respectively. Plant Direct 2, e00049.

Yang, J., Liu, Y., Yan, H., Tian, T., You, Q., Zhang, L., Xu, W., and Su, Z. (2018). PlantEAR: Functional analysis platform for plant EAR motif-containing proteins. Front. Genet. 9, 590.

Yang, T., Wu, X., Wang, W., and Wu, Y. (2023). Regulation of seed storage protein synthesis in monocot and dicot plants: A comparative review. Mol. Plant 16, 145–167.

Yin, L.L., and Xue, H.W. (2012). The MADS29 transcription factor regulates the degradation of the nucellus and the nucellar projection during rice seed development. Plant Cell 24, 1049–1065.

Zhang, L., Ren, Y., Lu, B., Yang, C., Feng, Z., Liu, Z., Chen, J., Ma, W., Wang, Y., Yu, X., Wang, Y., Zhang, W., Wang, Y., Liu, S., Wu, F., Zhang, X., Guo, X., Bao, Y., Jiang, L., and Wan, J. (2016). FLOURY ENDOSPERM7 encodes a regulator of starch synthesis and amyloplast development essential for peripheral endosperm development in rice. J. Exp. Bot. 67, 633–647.

Zhang, S., Dong, R., Wang, Y., Li, X., Ji, M., and Wang, X. (2021). NAC domain gene VvNAC26 interacts with VvMADS9 and influences seed and fruit development. Plant Physiol. Biochem 164, 63–72.

Zhang, Y., Chen, M., Siemiatkowska, B., Toleco, M.R., Jing, Y., Strotmann, V., Zhang, J., Stahl, Y., and Fernie, A.R. (2020). A highly efficient *Agrobacterium*-mediated method for transient gene expression and functional studies in multiple plant species. Plant Commun. 1, 100028.

Zhang, Y.-J., Zhang, Y., Zhang, L.-L., He, J.-X., Xue, H.-W., Wang, J.-W., and Lin, W.-H. (2022). The transcription factor OsGATA6 regulates rice heading date and grain number per panicle. J. Exp. Bot. 73, 6133–6149.

Zhang, Z., Dong, J., Ji, C., Wu, Y., and Messing, J. (2019). NAC-type transcription factors regulate accumulation of starch and protein in maize seeds. Proc. Natl. Acad. Sci. U S A 116, 11223–11228.

Zhao, X., Yang, X., Pei, S., He, G., Wang, X., Tang, Q., Jia, C., Lu, Y., Hu, R., and Zhou, G. (2016). The *Miscanthus* NAC transcription factor MlNAC9 enhances abiotic stress tolerance in transgenic *Arabidopsis*. Gene 586, 158–169.

Zhu, F., Luo, T., Liu, C., Wang, Y., Zheng, L., Xiao, X., Zhang, M., Yang, H., Yang, W., Xu, R., Zeng, Y., Ye, J., Xu, J., Xu, J., Larkin, R.M., Wang, P., Wen, W., Deng, X., Fernie, A.R., and Cheng, Y. (2020). A NAC transcription factor and its interaction protein hinder abscisic acid biosynthesis by synergistically repressing NCED5 in *Citrus reticulata*. J. Exp. Bot. 71, 3613–3625.

Zhuang, H., Wang, H.L., Zhang, T., Zeng, X.Q., Chen, H., Wang, Z.W., Zhang, J., Zheng, H., Tang, J., Ling, Y.H., Yang, Z.L., He, G.H., and Li, Y.F. (2020). NONSTOP GLUMES1 encodes a C2H2 zinc finger protein that regulates spikelet development in rice. Plant Cell 32, 392–413.

Zou, X., Neuman, D., and Shen, Q.J. (2008). Interactions of two transcriptional repressors and two transcriptional activators in modulating gibberellin signaling in aleurone cells. Plant Phys. 148, 176–186.

